# Integrating activation-induced costimulation and cytokine signals enhance TCR-based cell therapies

**DOI:** 10.64898/2026.07.12.738084

**Authors:** Mansi Narula, Johannes Englisch, Cheryl Ou, Tadahiro Honda, Ismael de la Iglesia-San Sebastian, Azlann Arnett, Feiyan Mo, Maksim Mamonkin, Norihiro Watanabe

**Affiliations:** Center for Cell and Gene Therapy, Baylor College of Medicine, Texas Children’s Hospital and Houston Methodist Hospital, Houston, Texas, USA; Graduate Program in Immunology and Microbiology, Baylor College of Medicine, Houston, TX 77030, USA; Center for Advanced Innate Cell Therapy, Texas Children’s Hospital, Houston, Texas, USA

## Abstract

TCR-based cell therapies offer broad targeting of tumor antigens with high sensitivity but often low durability due to insufficient costimulation (signal 2) and cytokine (signal 3) support. To address this, we developed a dual stimulatory receptor (DSR) consisting of the 4-1BBL ectodomain fused to the thrombopoietin receptor (cMPL) endodomain. DSR engages 4-1BB that is transiently upregulated upon TCR stimulation eliciting signal 2 and simultaneously activates cMPL-driven STAT3/5 phosphorylation providing signal 3. DSR increases the expansion of T cells, preserving their effector function and an effector-associated transcriptional profile upon repeated antigen stimulation. DSR arming significantly improves *in vivo* antitumor activity of T cells redirected to cancer through engineered TCRs or soluble T-cell engagers in diverse xenograft tumor models by enhancing T cell expansion and persistence post-infusion. These results demonstrate broad utility and establish DSR as a modular receptor for the effective and synchronized delivery of signals 2 and 3 to support TCR-based immunotherapies.

## Introduction

T cell receptor (TCR)-based T cell therapy has been the cornerstone of cellular immunotherapy for cancer^1^. Owing to their ability to recognize a broad repertoire of tumor antigens presented by the major histocompatibility complex (MHC) and to leverage physiological antigen sensitivity, there is a growing spectrum of TCR-based therapeutic modalities under investigation^2^. Early efforts focused on harnessing the potential of native TCRs through *ex vivo* expansion of virus-specific T cells (VSTs)^3,4^ and tumor-infiltrating lymphocytes (TILs)^5,6^. These approaches have since evolved towards the genetic engineering of TCRs (TCR-T) to confer defined specificity against tumor-associated antigens (TAAs) or neoantigens^7^. Furthermore, antibody-like Immunotherapeutics that recruit the endogenous TCR to the tumor antigen has led to the development of soluble T cell engagers (TCEs)^8^. The success of TCEs is evident by their widespread clinical use and several other iterations currently under development^9^. More recently, innovative approaches have been used to integrate the antigen specificity of single-chain variable fragment (scFv) with TCR/CD3 signaling machinery gave rise to synthetic chimeric TCRs^2,10^. These strategies have largely focused on enhancing antigen recognition efficiency which is sufficient to transmit the cytolytic signal to T cells (signal 1), but most do not integrate concomitant costimulatory signal (signal 2) and cytokine signal (signal 3), critical for a sustained anti-tumor response. In the absence of these signals, repeated T cell dosing and/or cytokine supplementation is necessary to achieve a durable *in vivo* efficacy^11–13^. Although approaches to incorporate such signals directly into the TCR complex have been investigated, their functional benefit was limited due to structural constraints^14–17^. On the other hand, *in trans* engineering strategies providing either signal 2 or signal 3 have been extensively explored. Arming T cells with synthetic cytokine signaling, such as the secreted IL-12^18^ or the constitutive IL-7 receptor (C7R)^19^, improve T cell expansion and/or their phenotype; however, such constitutive signaling raises concern regarding systemic toxicity or antigen-independent T cell expansion. Alternative approaches, including costimulatory CAR (coCAR)^20^, chimeric costimulatory receptors (CCRs)^21^, and synthetic cytokine circuits^22^, instead deliver orchestrated signals 2 or 3 triggered by secondary tumor antigens or the microenvironment, incurring dependency on factors extrinsic to T cells. Moreover, engineering antigen-specific T cells to express costimulatory ligands, including CD80 and 4-1BBL^23,24^, or dual-costimulatory variants (80BB)^25^, enables native signal 2 costimulation through the engagement of endogenous receptors (CD28 and 4-1BB). Therefore, most existing strategies engage either signals 2 or 3, but rarely both, particularly in the context of TCR-based therapies^26^.

In this study, we sought to address these limitations by developing a dual stimulatory receptor (DSR) that integrates costimulatory and cytokine signaling within a single synthetic construct. DSR is composed of a costimulatory ligand ectodomain fused to an immunostimulatory endodomain. The ectodomain is designed to engage with an endogenous (native) costimulatory receptor 4-1BB, induced on T cells upon antigen stimulation, to provide signal 2, while this interaction simultaneously activates the DSR endodomain to transmit signal 3. Through this design, both signals 2 and 3 are induced by T cell activation (signal 1) and independently of external factors such as tumor antigens or the tumor microenvironment (TME). Importantly, DSR-engineered T cells exhibited superior *in vivo* antitumor activity irrespective of the type of tumor antigen or targeting TCR, supporting the versatility of this approach for TCR-based immunotherapy.

## Results

### Dual stimulatory receptor (DSR) enhances T cell expansion upon repeated TCR stimulation

To create a dual stimulatory receptor that elicits both signals 2 and 3 upon TCR stimulation, we coupled the 4-1BBL ectodomain to the thrombopoietin receptor endodomain (cMPL) *via* a CD28 transmembrane region (**Figure 1a**). As the endogenous costimulatory receptor 4-1BB is transiently upregulated following T cell activation upon tumor recognition (signal 1), we hypothesized that its interaction with the 4-1BBL ectodomain of DSR would transmit signal 2 through endogenous 4-1BB and simultaneously trigger cytokine-like signal 3 through the cMPL endodomain of DSR, independent of extrinsic cues. Control constructs encoding a full-length wild-type 4-1BBL (4-1BBL WT) and a truncated DSR lacking the cMPL endodomain (ΔDSR) were generated to provide signal 2 through endogenous 4-1BB but no signal 3. Notably, since 4-1BBL WT is a type II transmembrane protein, the proximal-distal sequence of the ectodomain was reversed for a type I 4-1BBL-cMPL fusion DSR or ΔDSR construct (**Figure 1a**). Gene-modified T cells were generated in human platelet lysate (HPL)-supplemented medium to maintain a less-differentiated memory phenotype as previously shown^27^. All constructs were efficiently transduced into human T cells without affecting the cell expansion and memory phenotype (**Figure 1b** and Extended Figure 1a). While 4-1BBL WT and ΔDSR were detectable on the cell surface at rest, DSR was predominantly intracellular and expressed on the surface upon T cell activation (**Figure 1c** and **1d**). To evaluate the effect of DSR on T cell expansion upon TCR stimulation in the absence of exogenous cytokines, we repeatedly stimulated them using anti-CD3 scFv (OKT3)-expressing irradiated K562 cells. DSR-modified T cells continued to expand for at least six stimulation cycles, whereas control groups declined after 3 cycles (**Figure 1e**). Importantly, this enhanced cell expansion was stimulation-dependent, as antigen withdrawal resulted in rapid contraction of the DSR-modified T cells (**Figure 1f**).

**Figure 1:**
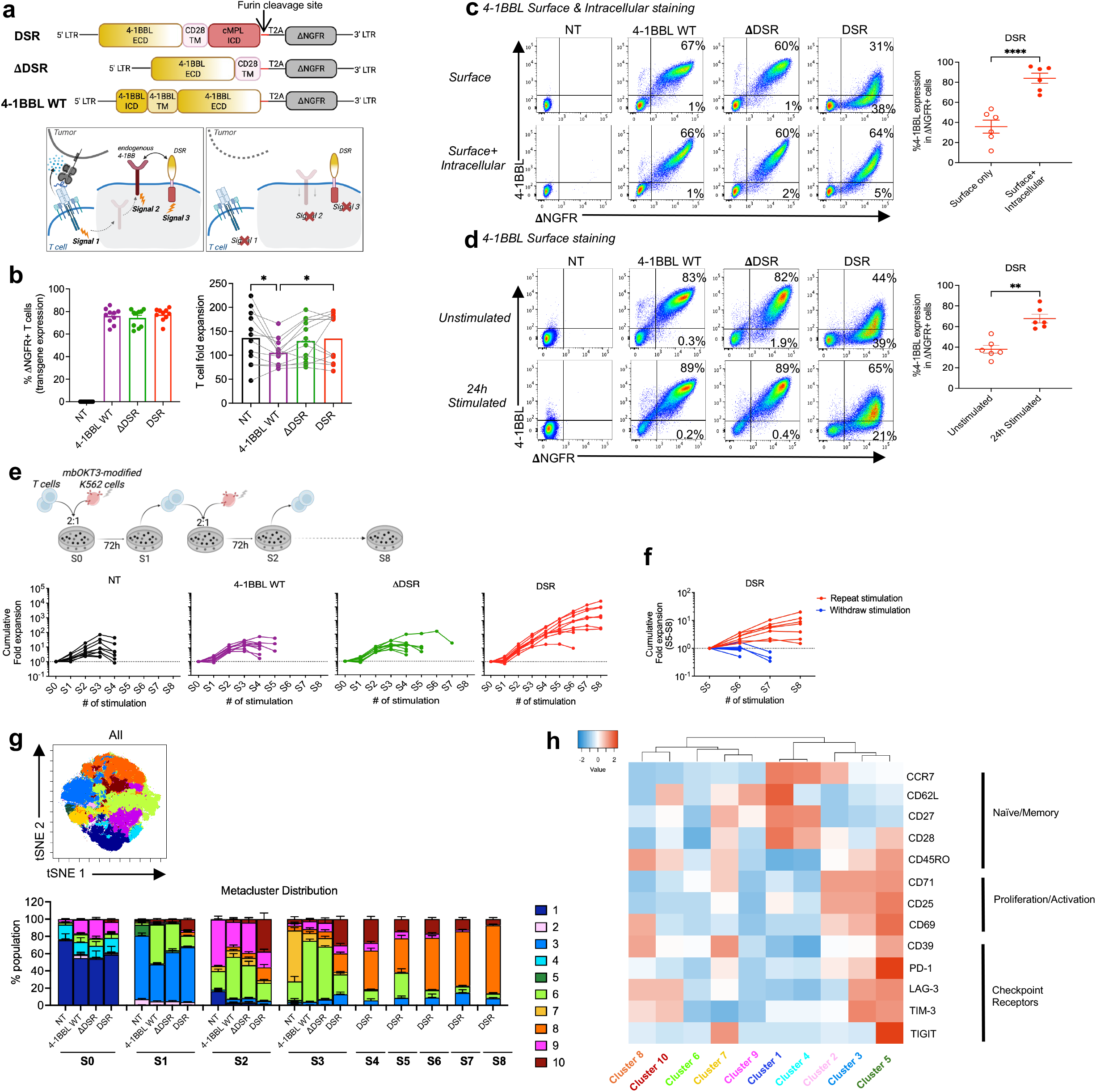
Dual stimulatory receptor (DSR) enhances T cell expansion upon repeated antigen stimulation. a, Bicistronic γ-retroviral construct map of DSR, ΔDSR and 4-1BBL WT with ΔNGFR surrogate marker (top) and expected mode of action of DSR-transduced T cells upon tumor recognition (bottom). **b,** ΔNGFR surface expression (n=10, left) and fold expansion (n=11, right) of gene-modified T cells (day 7 post transduction). **c,** Representative flow plots of 4-1BBL expression on the surface (top) and surface with intracellular staining (bottom) in resting state. Summary plot showing %4-1BBL(+)within ΔNGFR(+) DSR-modified T cells (right, n=6). **d,** Representative flow plots of surface 4-1BBL expression in unstimulated (top) and OKT3-stimulated (24 hrs, bottom) gene-modified T cells. Summary plot showing %4-1BBL(+) within ΔNGFR(+) DSR-modified T cells (n=6). **e,** Schematic of repeated stimulation assay and line graphs showing cumulative T cell expansion upon repeated stimulation (S0-S8) in each experimental group (n=9/group). **f,** Cumulative T cell expansion of DSR-modified T cells with (red) or without (blue) stimulation (n=5). **g**, Concatenated tSNE-CUDA plot with 10 color-coded FlowSOM metaclusters of T cell phenotype upon repeated stimulation (n=5, left). Stacked bar graph showing the metacluster distribution across each experimental group and stimulation cycle (S0-S8, right). **h,** z-score heatmap of surface marker expression levels (n=5) with hierarchical clustering across metaclusters. All data with multiple donors shown as Mean ± S.E. Statistical analysis was performed by RM one-way ANOVA with Tukey’s multiple comparison in panel b, paired t-test in panels c,d. ^✱^p ≤ 0.05, ^✱✱^p ≤ 0.01, ^✱✱✱^p ≤ 0.001, ^✱✱✱✱^p ≤ 0.0001. ECD: extracellular domain, ICD: Intracellular domain, TM: transmembrane, NT: non-transduced.

As we observed progressive enrichment of CD8(+) T cells upon repeated antigen stimulation (Extended Figure 1b), CD8(+) T cell phenotype was analyzed at the end of each stimulation cycle. t-distributed stochastic neighbor embedding (tSNE) analysis followed by FlowSOM unsupervised clustering revealed the distribution of phenotypic clusters across groups and stimulation cycles (**Figure 1g** and Extended Figure 1c). While all groups transitioned from a naïve-like state (metacluster 1) to an activated phenotype (metacluster 3) post 1^st^ stimulation (S1), divergence emerged after the 2^nd^ stimulation (S2-S8) (**Figure 1g** and **1h**). DSR-modified T cells preferentially accumulated a differentiated CD45RO(+) CCR7(-) population co-expressing LAG-3, TIM-3, CD69 and CD39 (metacluster 8), whereas 4-1BBL WT and ΔDSR groups displayed features of terminal differentiation (metacluster 6) preceding proliferative collapse. Overall, persisting DSR-modified T cells exhibited a distinct phenotype consistent with their sustained expansion *in vitro*.

### DSR-modified T cells retain the effector function and gene signature upon repeated stimulation

To assess whether sustained expansion was accompanied by preserved function, we evaluated cytokine secretion at early (S1_24h) and late (S4_24h) cycle during repeated antigen stimulation, as indicated in **Figure 2a**. While minimal cytokine production was detected without stimulation (Extended Figure 2a), all groups produced effector cytokines following the 1^st^ stimulation (S1_24h). Strikingly, only DSR-modified T cells retained robust cytokine secretion post 4^th^ stimulation (S4_24h, **Figure 2b**), suggesting functional fitness consistent with their superior expansion at this time point (**Figure 1e**). To further dissect functional differences, we performed transcriptomic profiling that revealed minimal differences among groups at early stimulation cycle (S0 and S1_24h) with substantial divergence observed at the later stimulation cycles (S3, S4_24h, **Figure 2c**). Notably, DSR-modified T cells at S4_24h maintained transcriptomic similarity to S1_24h (**Figure 2c** and Extended Figure 2b). DSR-modified T cells showed an increased number of differentially expressed genes (DEGs) against the other groups at S4_24h (DSR vs NT: 561, DSR vs 4-1BBL WT:133, DSR vs ΔDSR:163) than at S1_24h (DSR vs NT:100, DSR vs 4-1BBL WT:91, DSR vs ΔDSR:37), suggesting growing differences in the transcriptional program upon repeated stimulation (Extended Figure 2c). Additionally, we detected a convergent set of upregulated (MAF^28^, RCAN2^29^, SPOCK1^30^, ROR2^31^, FGFBP2^32^) and downregulated (CCL22^33^, JPH4^34^, COL1A1, NTRK2^35^) genes in DSR-modified T cells versus T cells expressing either 4-1BBL WT or ΔDSR. These genes are not reported to be directly regulated by c-MPL signaling, however, they are associated with either of the three distinct signaling cascades downstream of cMPL-JAK2, namely STAT3/5, MAPK and PI3K-Akt^36,37^.

**Figure 2:**
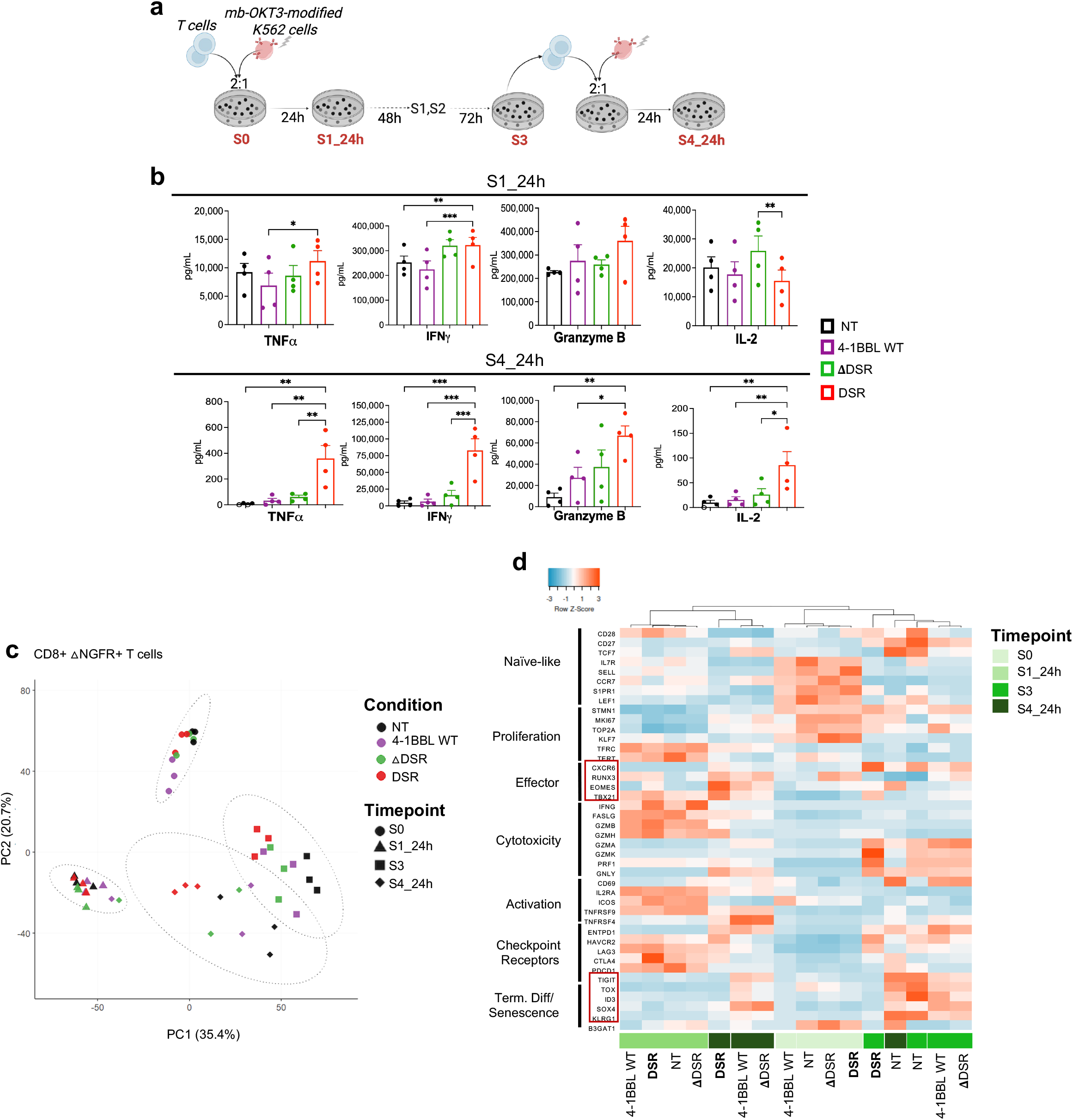
DSR-modified T cells maintain polyfunctionality and effector gene signature upon repeated antigen stimulation. **a**, Schematic of sample collection time points post repeated stimulation. **b,** Summary plots of effector cytokine secretion in T cells at indicated time points (Mean ± S.E., n=4). Open symbols indicate values below detectable limit **c,** Principal component analysis (PCA) of RNA-seq data from sorted ΔNGFR(+) CD8(+) T cells (n=3) in each experimental condition and time point specified in panel a. **d,** z-score heatmap of normalized gene expression (mean of 3 donors) of a selected gene set in each group and time point specified in panel a with hierarchical clustering across columns. Statistical analysis was performed by RM one-way ANOVA with Dunnett’s multiple comparison (vs DSR) in panel b. ^✱^p ≤ 0.05, ^✱✱^p ≤ 0.01, ^✱✱✱^p ≤ 0.001, ^✱✱✱✱^p ≤ 0.0001

Relative expression levels of a curated gene set representing prominent T cell functions^38,39^ demonstrated a naïve-like and proliferative gene signature at S0, consistent with the cell phenotype (**Figure 2d**). As expected, upregulation of activation-induced (IL2RA, ICOS, TNFRSF9, HAVCR2, LAG3, CTLA4, PDCD1) and cytotoxic genes (IFNG, FASLG, GZMB, GZMH) were observed at S1_24h across all groups. Importantly, DSR-modified T cells uniquely maintained an elevated levels of cytotoxic gene expression at S4_24h. At S3 and S4_24h, DSR-modified T cells exhibited higher effector-associated gene signature (TBX21, EOMES and CXCR6) and lower terminal differentiation and senescence-associated genes (TOX, ID3, SOX4, KLRG1), resulting in closer hierarchical clustering of S1_24h and S4_24h DSR-modified T cells, suggesting maintenance of effector transcriptomic profile (**Figure 2d**). Gene set enrichment analysis (GSEA) further revealed enrichment of pathways related to mitochondrial metabolism, intact cell-cycle progression, and TCR and inflammatory cytokine signaling in DSR-modified T cells at S4_24h (Extended Figure 2d). Together, these findings indicate that DSR preserves polyfunctionality and an effector-associated transcriptional program despite repeated antigen exposure.

### 4-1BB interacts with DSR to turn on critical T cell signals 2 and 3

To define how DSR produces functional benefit and an active gene signature in T cells, we visualized the localization and interaction between DSR and 4-1BB by utilizing fluorescence-tagged constructs (Extended Figure 3a). Imaging analysis revealed distinct localization patterns, with mEmerald-tagged 4-1BBL WT and ΔDSR at the cell surface, while mEmerald-DSR was predominantly intracellular (Extended Figure 3b), consistent with flow cytometry (**Figure 1c)**. When co-expressed with mScarlet-tagged Δ4-1BB, 4-1BBL WT and ΔDSR co-localized with Δ4-1BB *in cis* at the cell surface, whereas DSR and Δ4-1BB co-localized intracellularly (**Figure 3a**). Furthermore, flow cytometric analysis demonstrated the interaction between 4-1BBL and endogenous 4-1BB evidenced by the reduced detectable surface 4-1BB in 4-1BBL gene-modified T cells (**Figure 3b**). Comparable 4-1BB protein levels observed in NT, 4-1BBL WT and ΔDSR-modified T cells by western blot confirmed an effect of epitope masking by 4-1BBL contributing to low detection by flow cytometry (Extended Figure 3c). In contrast, relatively lower 4-1BB was observed in DSR-modified T cells by western blot, although 4-1BB mRNA levels remained unchanged among groups. This indicated post-translational differences and potential 4-1BB degradation^40^ rather than transcriptional changes in 4-1BB (Extended Figure 3d).

**Figure 3:**
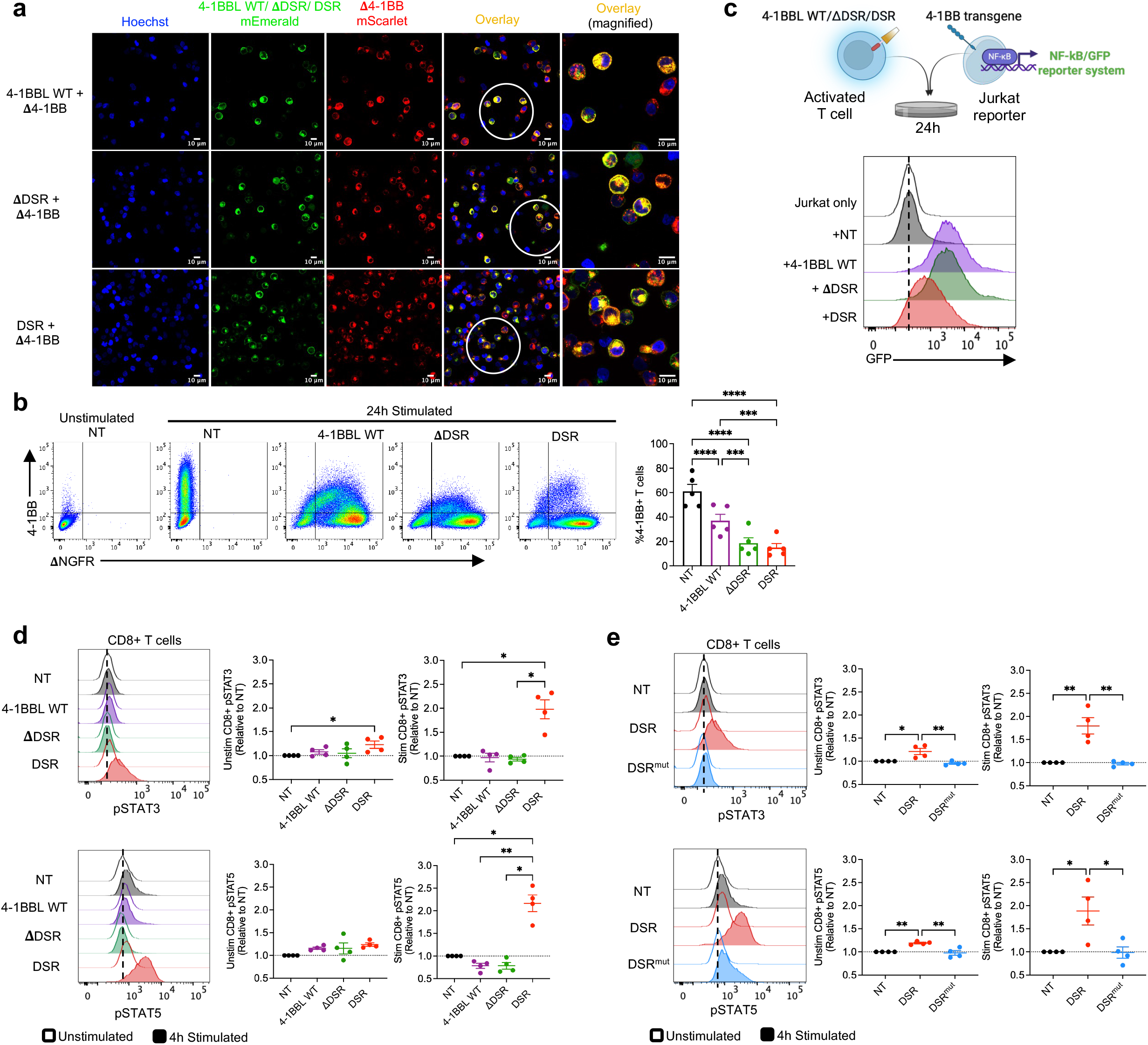
DSR engages 4-1BB and provides signals 2 and 3 to T cells. a, Fluorescence microscopy images of gene-modified T cells co-expressing mEmerald (green)-tagged 4-1BBL WT, ΔDSR or DSR and mScarlet (red)-tagged Δ4-1BB. Hoechst (blue) indicates nuclear stain and co-localization shown in yellow overlay. Images were acquired with a 63x objective lens (left) and 2.5x magnification zoom (right). Magnified view of the overlay field-of-view within the white circle. **b,** Representative flow plots of endogenous surface 4-1BB in T cells with or without OKT3-stimulation (left). Summary plots showing %4-1BB(+) T cells (n=5, right). **c,** Schematic of 4-1BB expressing NF-kB Jurkat reporter cell line cocultured with activated (for 24 hrs) gene-modified T cells at 1:1 (top). Histograms showing GFP induction in Jurkat cells at the end of coculture (bottom). **d,** Representative histograms of phospho-STAT3 and phospho-STAT5 staining in unstimulated (open) or 4 hr OKT3-stimulated (filled) CD8(+) T cells (left). Summary plots showing gMFI relative to NT cells (n=4, right). **e,** Representative histograms of phospho-STAT3 and phospho-STAT5 staining in unstimulated (open) or 4 hr OKT3-stimulated (filled) CD8(+) T cells modified with JAK2-binding site DSR mutant (DSR^mut^). Summary plots showing gMFI relative to NT cells (n=4, right). All data with multiple donors shown as Mean ± S.E. Statistical analysis was performed by RM one-way ANOVA with Tukey’s multiple comparison in panel b and Dunnett’s multiple comparison (vs DSR) in panels d,e. ^✱^p ≤ 0.05, ^✱✱^p ≤ 0.01, ^✱✱✱^p ≤ 0.001, ^✱✱✱✱^p ≤ 0.0001

To determine whether the 4-1BBL moiety in DSR can induce 4-1BB signaling, DSR-modified T cells, pre-activated to induce surface 4-1BBL expression, were co-cultured with 4-1BB expressing Jurkat NFκB-GFP reporter line (**Figure 3c** and Extended Figure 3e). GFP induction was observed with 4-1BBL WT or ΔDSR as well as with DSR-modified T cells, corresponding to the 4-1BBL surface expression levels, demonstrating the capacity of *in trans* 4-1BB activation even with the inverted 4-1BBL orientation in DSR (**Figure 3c**). Next, to assess cMPL endodomain signaling in DSR, we evaluated phosphorylation of STAT3 and STAT5^37,41^. While DSR-modified T cells demonstrated slightly higher background pSTAT3/5 relative to NT cells in the absence of stimulation, this effect was significantly enhanced upon T cell activation (**Figure 3d** and Extended Figure 3f). Notably, disruption of the cMPL Box 1 - JAK2 binding site^42,43^ (Extended Figure 3g) abrogated the STAT signal, confirming cMPL-dependent signaling in DSR (**Figure 3e** and Extended Figure 3h). Overall, the data validates DSR engagement with 4-1BB and induction of cMPL signaling, thereby providing coordinated signals 2 and 3.

### DSR boosts the *in vivo* persistence and anti-tumor function of transgenic TCR-modified therapeutic T cells

Finally, we evaluated the impact of DSR on the anti-tumor activity of therapeutic T cells across multiple modalities. First, we used a transgenic TCR specific to the survivin peptide/HLA-A2 complex, broadly expressed in multiple cancer types^12,44,45^ (**Figure 4a**). CD8(+) T cells were engineered to co-express survivin-specific TCR (SurvTCR) and DSR, where DSR expression did not alter the phenotype of SurvTCR-T cells (Extended Figure 4a and 4b). In a 9-day *in vitro* coculture with BV173 leukemia cells in the absence of exogenous cytokine supplementation, SurvTCR alone mediated only transient tumor control (Day 3), followed by the loss of T cell expansion and subsequent tumor outgrowth (**Figure 4b-d**). Co-expression with 4-1BBL WT or ΔDSR produced a modest improvement in T cell expansion and anti-tumor activity, whereas DSR enabled sustained T cell expansion and durable tumor control. This underscores the requirement for coordinated delivery of all three signals in TCR-engineered therapies.

**Figure 4:**
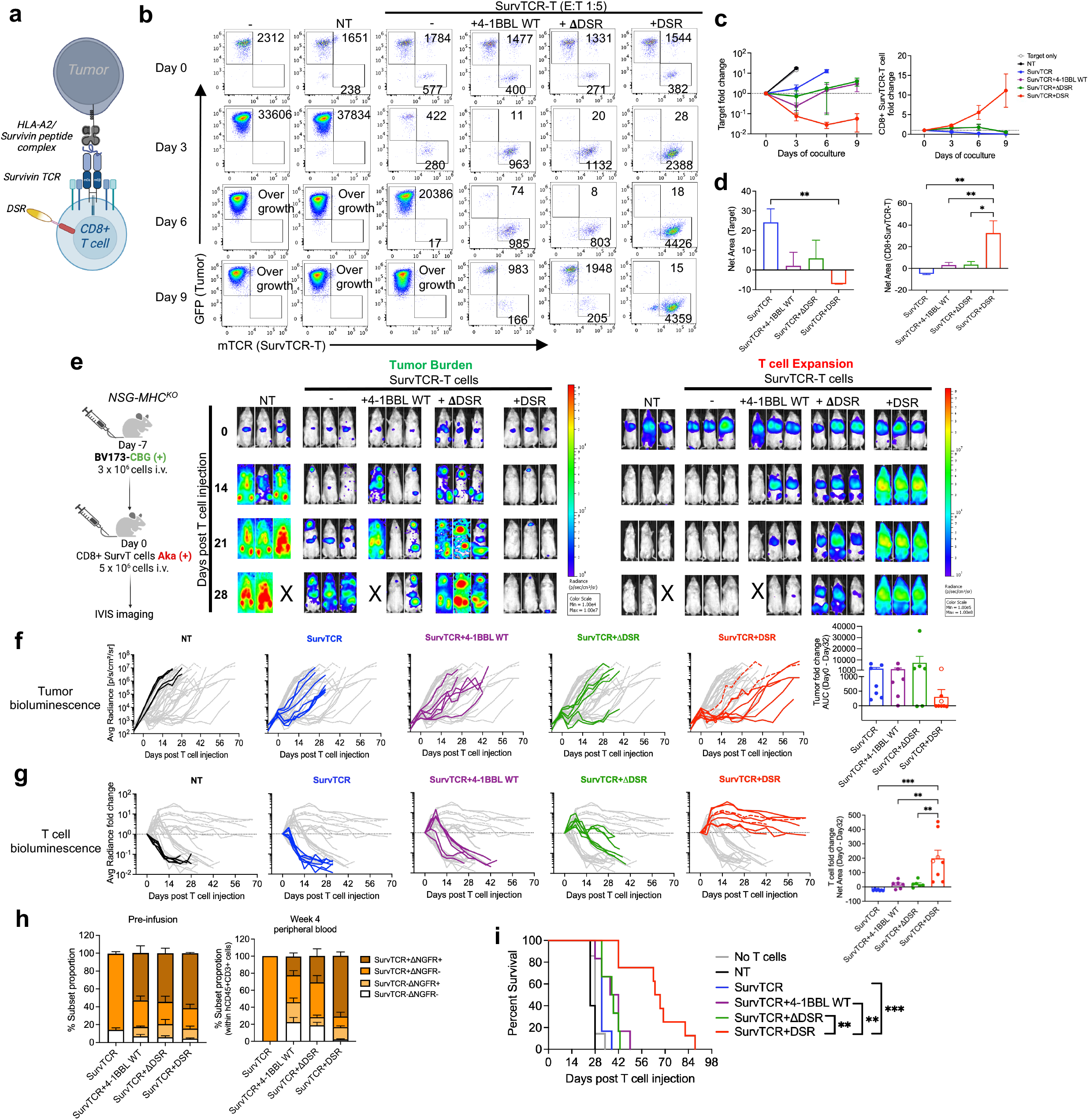
DSR promotes *in vivo* T cell expansion and persistence of Survivin-specific transgenic TCR-T cells. a, Schematic of CD8(+) T cells co-expressing transgenic Survivin TCR and DSR targeting the tumor. **b,** Representative flow plots of 9-day coculture experiments of gene-modified Survivin TCR-T cells with GFP(+) BV173 leukemia cells at 1:5 effector to target ratio. **c,** Summary line graphs of tumor growth (n=6, left) and Survivin TCR(+) T cell expansion kinetics (n=6, right). **d,** Summary bar graphs of the net area for tumor (left) and Survivin TCR(+) T cell (right) expansion over 9 days of coculture (baseline=1). **e,** Schematic of the (CBG+) BV173 xenograft tumor model with (Aka+) T cell treatment in a dual luciferase system (left). Representative bioluminescent images of the tumor (CBG: middle) and T cells (Aka: right) (NT; n=5, SurvTCR; n=7, SurvTCR+4-1BBL WT; n=6, SurvTCR+ΔDSR; n=6, SurvTCR+DSR; n=8). **f,** Line graphs showing tumor bioluminescence over time (left) and bar graph showing the area-under-the-curve (AUC) over the first 32 days post T cell treatment in mice (right). Dotted lines and open circles indicate mice euthanized due to extramedullary tumor growth. **g,** Line graphs showing fold change of T cell bioluminescence (left) and bar graph showing net area for T cell expansion over the first 32 days post T cell treatment in mice (right, baseline=1). Dotted lines and open circles indicate mice euthanized due to extramedullary tumor growth. **h,** Stacked bar graphs showing Survivin TCR(+) and ΔNGFR(+) (as a surrogate marker for 4-1BBL WT, ΔDSR, DSR) T cells in the infusion product (left, n=1,total 3 batches) and within human T cells in the mice peripheral blood post-infusion (right, SurvTCR; n=1, SurvTCR+4-1BBL WT; n=3, SurvTCR+ΔDSR; n=4, SurvTCR+DSR; n=8). **i,** Overall mice survival post T cell treatment. All data with multiple donors shown as Mean ± S.E. Statistical analysis was performed by RM one-way ANOVA with Tukey’s multiple comparison in panels d,f,g and survival analysis was performed by Log rank (Mantel-Cox) test in panel i. ^✱^p ≤ 0.05, ^✱✱^p ≤ 0.01, ^✱✱✱^p ≤ 0.001, ^✱✱✱✱^p ≤ 0.0001

Encouraged by these findings, we assessed the DSR function *in vivo* using NSG-MHC^KO^ mice, to reduce the risk of xenogeneic graft-versus-host disease. Mice were engrafted with Click Beetle Green (CBG)*-*modified BV173 leukemia cells followed by a single injection of freshly thawed Aka Luciferase (Aka)-modified CD8(+) SurvTCR-T cells without exogenous cytokine injection (**Figure 4e**). Tumor growth, T cell expansion, and their localization were monitored by dual-luciferase *in vivo* imaging system (IVIS). Typical BV173 localization was observed in the bone marrow, lymph nodes, and liver, followed by migration and expansion of the infused T cells (**Figure 4e**). Consistent with *in vitro* results, SurvTCR-T cells alone showed only transient tumor control due to insufficient T cell expansion (**Figure 4f and 4g**). Addition of 4-1BBL WT or ΔDSR induced a short-term T cell expansion that peaked on day 7, but failed to support long-term persistence, resulting in limited improvement in the anti-tumor effect over SurvTCR alone. Notably, DSR-engineered SurvTCR-T cells exhibited robust expansion, prolonged persistence and delayed tumor progression (**Figure 4f and 4g**). Circulating human T cells showed selective enrichment of the antigen-specific DSR-expressing population, indicating preferential maintenance of functional tumor-reactive T cells *in vivo* (**Figure 4h**). This was reflected in significantly prolonged mice survival (**Figure 4i**). Although tumor emergence at extramedullary sites, previously observed in the BV173 xenograft model^46^, was not controlled by circulating SurvTCR(+) DSR(+) T cells (**Figure 4f** and Extended Figure 4c), DSR nonetheless conferred clear functional benefits to transgenic TCR-modified T cell therapy.

### DSR-engineered bispecific T-cell engager secreting T cells (TCE-T) mediate long-term tumor eradication *via* native TCR targeting

To determine whether the benefit of DSR extended beyond transgenic TCRs, we evaluated its effect in bispecific T-cell engager-secreting T cells (TCE-T), which rely on native TCR signaling for cytotoxicity and typically lack costimulatory and cytokine support. This limits their durability in the absence of repeated administration of the engineered-T cells and/or exogenous cytokines^13,47^. We tested if DSR could overcome these limitations using CD123-targeting TCE-T cells, previously developed for acute myeloid leukemia (AML) treatment^13^ (**Figure 5a** & Extended Figure 5a). Co-expression of DSR (or controls, c-myc as the surrogate marker) did not alter the CD4(+)/CD8(+) distribution or the memory phenotype of TCE-T cells (Extended figure 5b-d). *In vitro*, CD123-targeting TCE-T cells showed limited T cell expansion and minimal control of MOLM-13 AML cells (**Figure 5b and 5c**). Additional expression of 4-1BBL WT or ΔDSR failed to enhance the anti-tumor activity. In contrast, DSR-modified TCE-T cells showed markedly increased T cell expansion, leading to tumor clearance (**Figure 5b** and **5c**).

**Figure 5:**
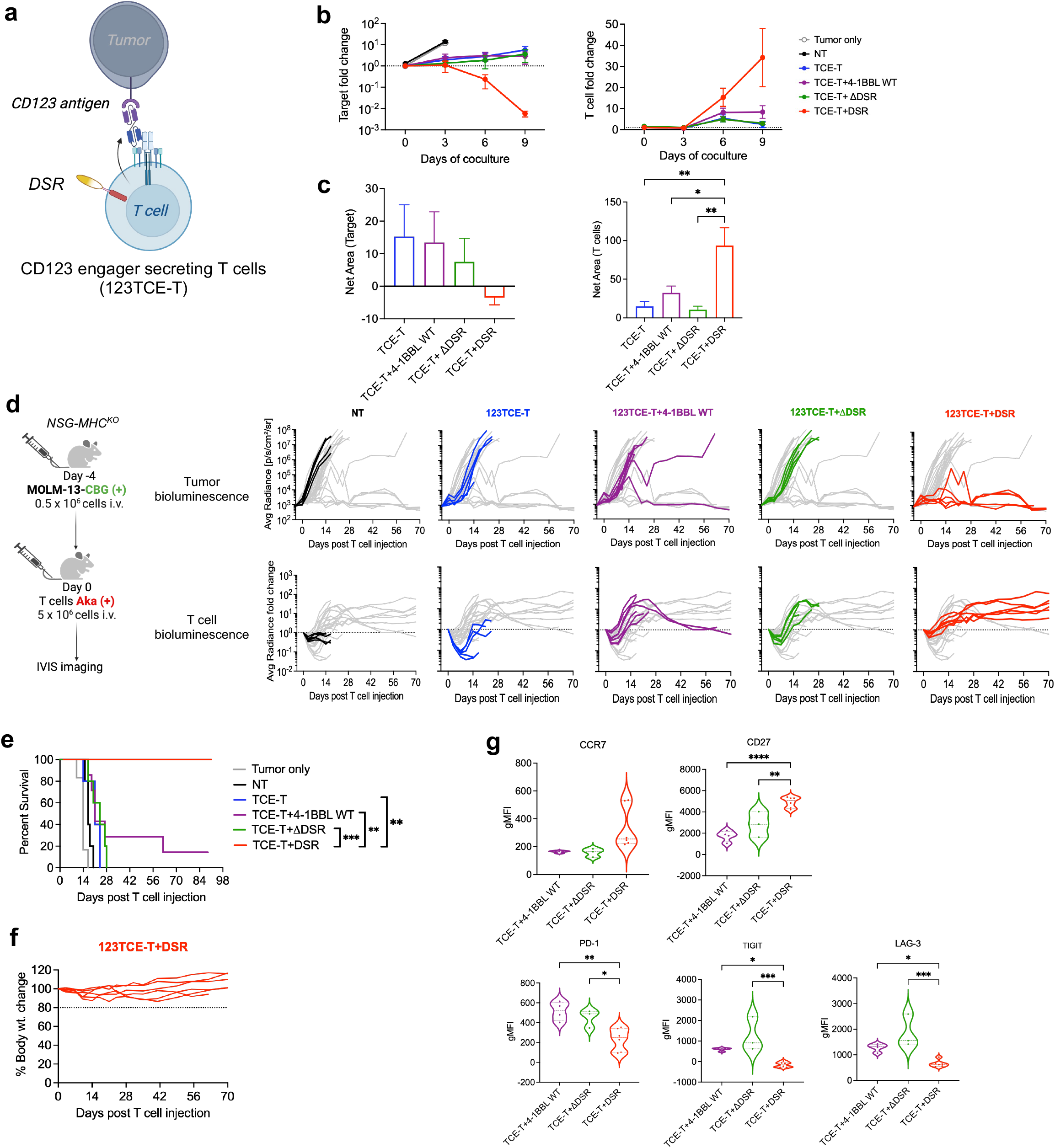
DSR-engineering of bispecific T cell engager-secreting T cells (TCE-T) enhances T cell persistence mediating long-term Acute Myeloid Leukemia (AML) control. a, Schematic of T cells secreting CD123-targeting engager molecule (123TCE-T) co-expressing DSR against AML cells. **b,** Summary line graphs of tumor growth (n=6, left) and T cell expansion kinetics (n=6, right) in a 9-day *in vitro* coculture of gene-modified 123TCE-T cells with GFP(+) MOLM-13 AML cell line at 1:4 effector to target ratio. **c,** Summary bar graphs of the net area for tumor (left) and T cell (right) expansion over 9 days of the coculture (baseline=1). **d,** Schematic of the CBG(+) MOLM-13 xenograft mouse model with Aka(+) T cell treatment in a dual-luciferase system (left). Line graphs showing tumor bioluminescence (CBG: top) and fold change in T cell bioluminescence (Aka: bottom) in mice per group (NT; n=5, 123TCE-T; n=5, 123TCE-T+4-1BBL WT; n=7, 123TCE-T+ΔDSR; n=5, 123TCE-T+DSR; n=6). **e,** Overall mice survival post T cell treatment. **f,** Line graphs showing % change in body weight over time relative to day 0 for each mouse treated with DSR-modified TCE-T cells (n=6). **g,** Violin plots summarizing the gMFI of each surface marker within the human CD8(+) T cells in the mice peripheral blood at 2 weeks post T cell infusion (TCE-T+4-1BBL WT; n=4, TCE-T+ΔDSR; n=3, TCE-T+DSR; n=6). All data with multiple donors shown as Mean ± S.E. Statistical analysis was performed by RM one-way ANOVA with Tukey’s multiple comparison in panel c, Ordinary one-way ANOVA with Dunnett’s multiple comparison (vs DSR) in panel g and survival analysis was performed by Log rank (Mantel-Cox) test in panel e. ^✱^p ≤ 0.05, ^✱✱^p ≤ 0.01, ^✱✱✱^p ≤ 0.001, ^✱✱✱✱^p ≤ 0.0001

In an aggressive MOLM-13 leukemia xenograft model injected with freshly thawed Aka (+) T cells (∼70% TCE-T, Extended Figure 5e), TCE-T cells alone or in combination with 4-1BBL WT or ΔDSR failed to control the tumor. In contrast, a single injection of DSR-engineered TCE-T cells without cytokine supplementation demonstrated robust T cell expansion and long-term persistence (**Figure 5d**). This resulted in durable tumor clearance and a significantly extended animal survival over ∼ 90 days (**Figure 5d and 5e**). Homeostatic levels of T cells persisting post tumor clearance was possibly an effect of the low-level DSR tonic cytokine signal as observed by the background STAT phosphorylation, previously in **Figure 3d**. Despite that, treated mice maintained stable body weight over the course of the study, indicating favorable tolerability with no systemic toxicity of DSR-modified T cells (**Figure 5f**). Notably, persisting DSR-modified T cells showed a favorable memory-associated (CCR7, CD27) and a less exhausted phenotype (PD-1, TIGIT, LAG-3) compared to 4-1BBL WT and ΔDSR groups (**Figure 5g**), despite higher peak T cell expansion with 4-1BBL WT or ΔDSR (Day 21, **Figure 5d**). Overall, the data depicts that DSR significantly benefits TCE-T cell native TCR-therapy and is effective, independent of the antigen-targeting modality.

### DSR enhances *in vivo* anti-tumor response of chimeric TCR-engineered T cells

CAR-TCR hybrid receptors that harness the antibody-like affinity of CARs (scFv or V_H_/V_L_) coupled to the TCR/CD3 signaling machinery represents a promising, newer strategy for targeting antigen-low tumors^2^. Although these novel chimeric receptors leverage native TCR signaling to deliver signal 1, they still lack signals 2 and 3. To evaluate whether DSR enhances the anti-tumor activity of this platform, we developed a CD19-specific chimeric TCR (19cTCR) consisting of a CD19-binding scFv fused to the mouse TCRα constant region and paired with mouse TCRβ1 constant region with specific mutations to improve cTCR stability^48,49^ (**Figure 6a** and Extended Figure 6a). Co-expression of DSR did not alter the characteristics of cTCR T cell products, similar to the previous two therapeutic T cell modalities (Extended Figure 6b-d). Functionally, DSR-modified 19cTCR-T cells cocultured with CD19(+) NALM6 cells exhibited enhanced and sustained proliferation, resulting in complete tumor elimination (**Figure 6b** and **6c**). This DSR-mediated benefit was attenuated upon endogenous 4-1BB gene knock-out, confirming the requirement for 4-1BB engagement to trigger DSR activity (**Figure 6d and 6e**, Extended Figure 6e).

**Figure 6:**
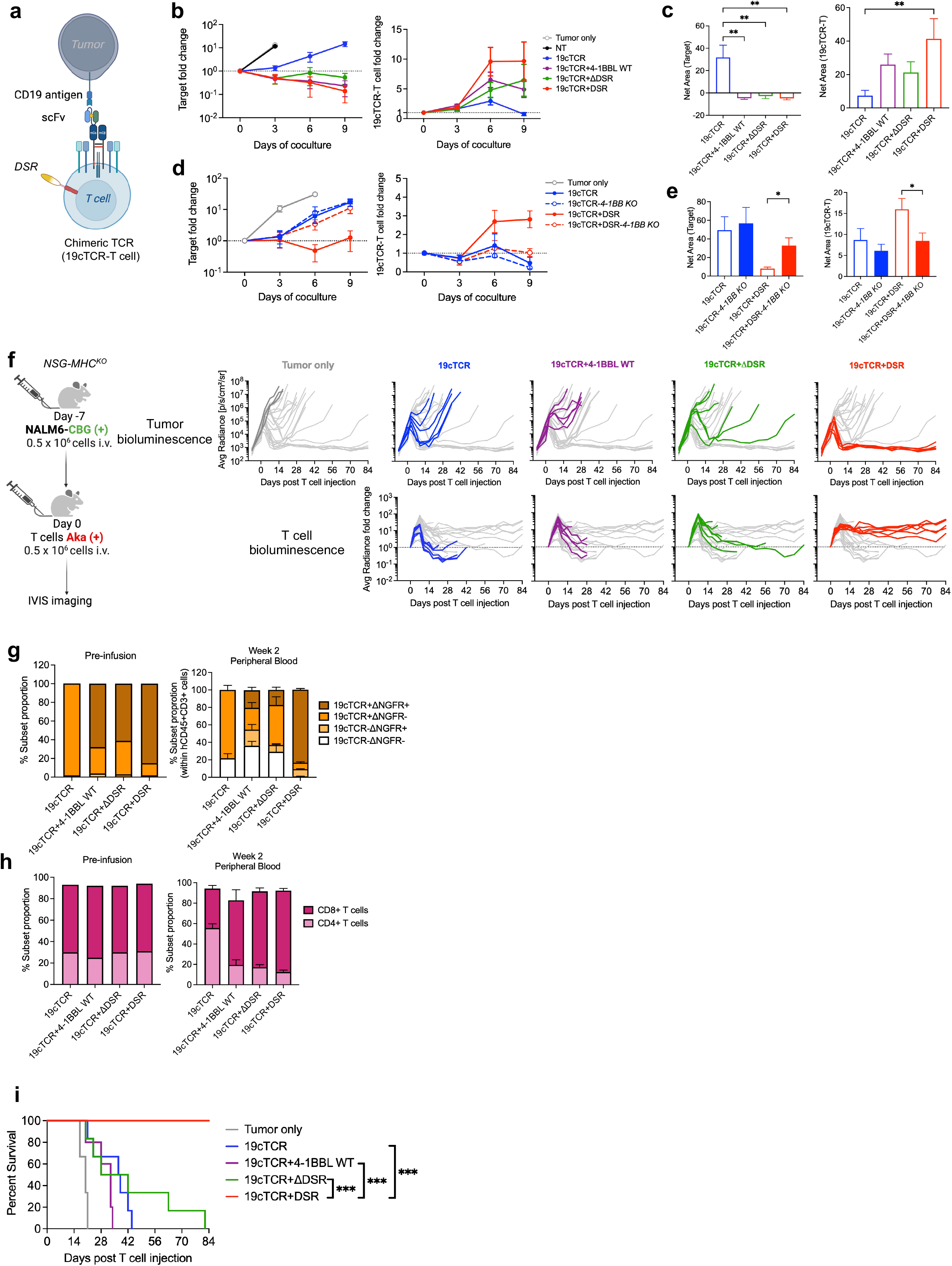
DSR enhances the T cell persistence and overall efficacy of CD19-chimeric TCR (19cTCR) targeting NALM6 Leukemia. a, Schematic of T cells co-expressing 19cTCR and DSR targeting leukemia cells. **b,** Summary line graphs showing tumor growth (n=6, left) and T cell expansion kinetics (n=6, right) in a 9-day *in vitro* coculture of gene-modified 19cTCR-T cells with GFP(+) NALM6 tumor at 1:5 effector to target ratio. **c,** Summary bar graphs of the net area for tumor (left) and T cell (right) expansion over 9 days of the coculture assay (baseline=1). **d,** Summary line graphs showing tumor growth (n=5, left) and T cell expansion kinetics (n=5, right) in a 9-day *in vitro* coculture of 19cTCR-T cells with or without endogenous 4-1BB knock-out with GFP(+) NALM6 tumor at 1:4 effector to target ratio. **e,** Summary bar graphs of the net area for tumor (left) and T cell (right) expansion in the 4-1BB knock-out coculture assay (baseline=1).**f,** Schematic of the CBG(+) NALM6 xenograft mouse model with Aka(+) T cell treatment in a dual luciferase system (left). Line graphs showing tumor bioluminescence (CBG: top) and fold change in T cell bioluminescence (Aka: bottom) in mice per group (No treatment; n=3, 19cTCR; n=6, 19cTCR+4-1BBL WT; n=6, 19cTCR+ΔDSR; n=6, 19cTCR+DSR; n=6). **g,** Stacked bar graphs showing 19cTCR(+) and ΔNGFR(+) (as a surrogate marker for 4-1BBL WT+/ΔDSR+/DSR+) in the infusion product (left) and within human T cells in the mice peripheral blood (right, 19cTCR; n=6, 19cTCR+4-1BBL WT; n=6, 19cTCR+ΔDSR; n=5, 19cTCR+DSR; n=6). **h,** CD4(+)/CD8(+) T cell distribution in the infusion product (left) and within human T cells in the mice peripheral blood (right, 19cTCR; n=6, 19cTCR+4-1BBL WT; n=6, 19cTCR+ΔDSR; n=5, 19cTCR+DSR; n=6). **i,** Overall mice survival post T cell treatment. All data with multiple donors shown as Mean ± S.E. Statistical analysis was performed by RM one-way ANOVA with Tukey’s multiple comparison in panel c, paired-t test in panel e, and survival analysis was performed by Log rank (Mantel-Cox) test in panel i. ^✱^p ≤ 0.05, ^✱✱^p ≤ 0.01, ^✱✱✱^p ≤ 0.001, ^✱✱✱✱^p ≤ 0.0001

In a CBG(+) NALM6 engrafted xenograft model, a single infusion of Aka(+) 19cTCR-T cells mediated only transient T cell expansion and short-term tumor control, with relapse occurring by day 10 post treatment (**Figure 6f**). Addition of 4-1BBL WT or ΔDSR increased peak T cell expansion but failed to provide long-term anti-tumor effect. In contrast, DSR-engineered 19cTCR-T cells exhibited sustained persistence and durable anti-tumor responses. As in other models, a selective enrichment of the DSR-modified tumor-reactive T cell fraction was observed in the peripheral blood, compared to a loss of tumor-reactive 4-1BBL WT or ΔDSR-modified T cells (**Figure 6f and 6g**). Consistent with the *in vitro* repeated stimulation assay (Extended Figure 1b), DSR modification also enriched the human CD8(+) T cell fraction in mice peripheral blood (**Figure 6h**). Overall, DSR-modified 19cTCR-T cells significantly prolonged animal survival over ∼ 80 days (**Figure 6i**). Together, our data emphasizes the versatility of DSR in producing a consistent effect across different TCR-based cell therapy platforms in diverse tumor models.

## Discussion

In this study, we describe a novel synthetic receptor, DSR, that simultaneously provides a costimulatory signal 2 and cytokine-like signal 3 to overcome an inherent limitation of TCR-based immunotherapies. We demonstrated that DSR signaling is initiated upon antigen stimulation (signal 1), inducing the expression of endogenous 4-1BB in activated T cells^50^ - the cognate binding partner of DSR. Through this mechanism, DSR enabled T cells to elicit signals 2 and 3 independently from the milieu. These properties distinguish DSR from previously established constructs that deliver either only signal 2 or signal 3, constitutively or through a secondary antigen trigger. By aligning signal 2 and 3 with antigen-driven signal 1, DSR reproducibly promoted robust yet controlled expansion and durable persistence of engineered T cells across different emerging therapeutic platforms.

Expression of full-length costimulatory ligands such as CD80 and 4-1BBL in T cells^23,24,46^, as well as a chimeric dual co-stimulatory 80BB receptor^25^, has been shown to deliver signal 2 through engagement of their cognate endogenous costimulatory receptors. On the other hand, ectopic expression of cMPL on T cells - normally restricted to hematopoietic stem cells and absent in mature T cells - elicits signaling analogous to common γ-chain cytokines in response to TPO, thereby enhancing T cell proliferation and anti-tumor activity^51^. Within the DSR framework, we delineated the discrete contributions of 4-1BBL and cMPL domains resulting in synergistic functional enhancement. Upon repeated stimulation, transcriptomic profiling of DSR-engineered T cells revealed upregulation of an inflammatory gene signature, including IFIT3, RSAD2, TBX21 and TNF, compared to unmodified T cells. This pattern aligns with prior studies describing a type I interferon-associated transcriptional program in T cells modified with either 4-1BBL^24^ or cMPL^51^.

Structurally, wild-type 4-1BBL naturally forms and exists as a stable trimer^52^ while ligand-dependent cMPL dimeric conformation leads to receptor internalization mediated through multiple cytoplasmic internalization sites^42,53^. Based on these premises, it is conceivable that DSR multimerization, driven by the 4-1BBL trimeric conformation, facilitates receptor internalization and reduced surface expression in resting state. This may also be contributing to the observed low-level antigen-independent tonic cytokine signal maintaining homeostatic levels of DSR-modified T cells in mice. In general, we observed accumulation of CD8(+) T cells across all groups upon *in vitro* repeated stimulation and this pattern was also preserved for DSR-engineered T cells in the *in vivo* models. This is consistent with previous reports linking 4-1BB signaling to enhanced CD8(+) T cell proliferation ^24,54^ and may reflect the prolonged expression kinetics of endogenous 4-1BB in this compartment^50^. Engagement of cMPL may further contribute to this skewness as seen with cMPL-embedded biosensor-modified CAR-T cells^37^.

Historically, the majority of TCR-based clinical trials have relied on exogenous cytokine supplementation, particularly with IL-2, to support T cell persistence^11,26^. Current efforts are increasingly focused on engineering antigen-reactive T cells, such as TILs, with complementary costimulatory signals (e.g. CD28 and CD40) or cytokine support (e.g. IL-7, IL-12 or IL-15) and evaluating them in clinical trials^55,56^. Although the clinical outcome reports from these approaches remain limited, early results from the Synthetic TCR and Antigen Receptor (STAR) construct incorporating OX40 costimulation demonstrate encouraging efficacy with acceptable safety in Phase I study of relapsed or refractory B cell leukemia^57^. On the other hand, multiple clinical trials have been launched to augment T cell engagers with costimulatory bispecifics^58^. In this context, DSR represents a further enhancement by integrating both costimulatory (signal 2) and cytokine-like (signal 3) inputs in concert with antigen-derived signal 1. From a translational perspective, DSR offers several advantages relevant to clinical applications. First, both the ectodomain and endodomain of DSR individually are known to function in a homomeric assembly, permitting delivery of DSR as a monomeric unit capable of multimerization, thereby enabling incorporation into compact, standard viral vectors. Second, self-sufficiency of T cells with DSR obviates the need for cytokine supplementation upon infusion. Finally, the compatibility of DSR across diverse TCR-based modalities examined in this study, including Survivin-specific TCR-transgenic T cells, CD123-redirected bispecific TCE-T cells, and CD19-scFv-containing chimeric TCR-T cells, highlight the breadth of DSR applications and the possibility of integration into existing clinical manufacturing workflows. Beyond conventional αβT cells, alternative cell types including invariant NKT cells^59^, γδT cells^60^, and natural killer cells^50,61^ also exhibit activation-induced upregulation of 4-1BB, with substantial evidence supporting the functional benefit of 4-1BB signaling in a therapeutic setting^62–64^. Given that insufficient cell persistence remains a common bottleneck across cell-based immunotherapies, DSR has the potential to improve the durability of a wide range of therapeutic platforms.

## Materials and Methods

### Donors and cell lines

Peripheral blood mononuclear cells (PBMCs) were isolated from whole blood obtained from healthy volunteers, with informed consent (BCM Protocol: H-45017). 293T (human embryonic kidney cell line), K562 (chronic myelogenous leukemia cell line), Jurkat (acute T-cell leukemia cell line), and NALM6 (pre-B-ALL cell line) were obtained from the American Type Culture Collection (Rockville, MD). BV173 (B-cell precursor leukemia cell line) and MOLM-13 (acute myeloid leukemia cell line) were obtained from the Leibniz Institute DSMZ-German Collection of Microorganisms and Cell Cultures GmbH (Braunschweig, Germany). 293T cells were maintained in DMEM (Dulbecco’s Modified Eagle Medium: Gibco BRL Life Technologies, Inc., Gaithersburg, MD) supplemented with 10% heat-inactivated fetal bovine serum (FBS) (Gibco BRL Life Technologies, Inc.) and 2 mM L-GlutaMAX (Gibco BRL Life Technologies, Inc.). K562, Jurkat, NALM6, MOLM-13, and BV173 cells were maintained in RPMI-1640 (HyClone Laboratories, Marlborough, MA) supplemented with 10% FBS and 2 mM L-GlutaMAX. All cells were maintained in a humidified atmosphere containing 5% carbon dioxide (CO_2_) at 37°C and were routinely tested for Mycoplasma.

### Generation of viral vectors and virus production

To generate the 4-1BBL WT bicistronic γ-retroviral vector (SFG), the full-length 4-1BBL fragment was gene-synthesized and cloned into the SFG backbone encoding a furin cleavage site (RAKR) and T2A sequence, followed by the ΔNGFR surrogate marker, using the NEBuilder HiFi DNA Assembly kit (New England Biolabs, Ipswich, MA). To generate DSR and ΔDSR constructs, the 4-1BBL ectodomain was PCR-amplified from 4-1BBL WT and assembled with the CD28 transmembrane (TM) region in the reverse orientation, with or without cMPL endodomain (gene synthesized), into the SFG backbone containing RAKR-T2A and ΔNGFR. We also generated the above constructs with a human c-myc tag (EQKLISEEDL) anchored on the membrane through a CD8 stalk and TM region in place of ΔNGFR. For fluorescence microscopy experiments, DSR (and 4-1BBL WT or ΔDSR controls) sequences were fused to the mEmerald sequence^50^ through a G4S linker and cloned into the SFG vector encoding a membrane-anchored FLAG-tag with a CD8 stalk and TM region as a surrogate marker. Similarly, the Δ4-1BB-mScarlet construct was cloned into the SFG vector encoding ΔNGFR by assembling the 4-1BB receptor ectodomain and TM region (PCR-amplified from the cDNA of activated T cells), lacking its endodomain to avoid toxicity due to continuous and excessive TNFR signaling^65^, with mScarlet^66^ via a G4S linker. In addition, a construct encoding the full-length 4-1BB into the SFG vector was also generated. The DSR-Box 1 mutant was generated by substituting the proline residues at position 17 and 20 within the Box 1 motif of the cMPL endodomain with alanine via site-directed mutagenesis^42^. To generate the lentiviral vector encoding the NFkB-GFP reporter, the sequence encoding the NFkB response element driving GFP was PCR-amplified from pSIRV-NF-kB-eGFP (Addgene: #118093), with replacement of the promoter region to a minimal TATA-box promoter, and cloned into a lentiviral vector encoding Q8 (CD34 epitope tag linked by a G_4_S linker to the CD8 stalk/transmembrane domain) as a surrogate marker. To generate the ΔFas construct, a Fas gene containing the signal peptide, extracellular domain, and transmembrane domain was PCR-amplified from cDNA of activated T cells and cloned into the SFG vector encoding ΔNGFR. The γ-retroviral vector encoding the survivin-specific TCR was previously published^12^ and kindly gifted by Dr. Bilal Omer (Baylor College of Medicine). The γ-retroviral vectors encoding the click beetle green luciferase co-expressing GFP (CBG/GFP), and Aka luciferase co-expressing GFP (Aka/GFP) were kindly gifted by Dr. Pradip Bajgain (NCI)^67^. The Aka/GFP vector was modified by replacing GFP with ΔEGFR, PCR-amplified from a DNA template provided by Dr. David Quach (Baylor College of Medicine). The γ-retroviral vector encoding the CD123 TCE with an mOrange surrogate marker was kindly gifted by Dr. Challice L. Bonifant (Johns Hopkins University)^13^ and modified to replace mOrange with ΔNGFR. To generate the γ-retroviral vector for CD19-cTCR, each fragment was assembled into SFG vector as follow: CD19 scFv-FMC63 with a FLAG tag (PCR-amplified from the CD19 CAR construct^62^, mouse Trac with mutations^48,49^ (gene synthesized), the furin-P2A sequence, and a myc-tagged mouse Trbc1 with mutation^49^ (gene synthesized). Primers were designed using SnapGene 6.0.5 software. Lentiviral and γ-retroviral supernatants were generated as previously described^65,68^.

### Generation of gene-modified T cells and cell lines

Gene-modified T cells were generated using our previously published protocol, with minor modifications^27,68^. Briefly, 10^6^ cells/mL PBMCs or purified CD8(+) T cells (for the generation of Survivin TCR-modified T cells), isolated with CD8(+) microbeads (Miltenyi Biotech Inc, San Diego, CA), were stimulated with plate-bound OKT3 (1 µg/mL) and anti-CD28 (1 µg/mL) antibodies in the presence of recombinant human IL-7 (10 ng/mL) and IL-15 (10 ng/mL) in CTL medium composed of 47.5% RPMI-1640, 47.5% Clicks medium (FUJIFILM Biosciences, Santa Ana, CA), and 2 mM L-GlutaMAX, supplemented with 5% of human platelet lysate (HPL; nLiven PR™, Sexton Biotechnologies, BioLife Solutions Inc., Bothell, WA). Two days later, activated T cells were retrovirally transduced on non-tissue culture 24-well plates coated with Retronectin reagent (Takara Bio USA, Inc., San Jose, CA) as previously described^68^. Cells were split and fed every 2– 3 days with fresh CTL medium plus IL-7/15 cytokines (10 ng/mL). Cells were harvested on day 7 post-transduction and cryopreserved. For *in vivo* experiments, Aka/ΔEGFR(+) cells were isolated using a biotin-conjugated anti-EGFR monoclonal antibody followed by anti-biotin microbeads (Miltenyi Biotec Inc.) using the MACS system during the manufacturing process. To generate cell lines overexpressing transgenes, we used the same protocol described above and isolated the transduced population using either a cell sorter (SH800S, Sony Biotechnology, San Jose, CA) or the MACS system. While gene-modified T cells were generated in CTL medium supplemented with 5% HPL, all *in vitro* functional assays were performed in CTL medium supplemented with 10% FBS.

### Gene knock-out (KO) in T cells and testing KO efficiency

A single-guide RNA (sgRNA) specific for 4-1BB (target sequence: GGGCTGGAGAAACTATTTGG) was designed using CRISPRscan and COSMID algorithms. sgRNA was generated as previously described^69^. For gene knock-out, 1 ug sgRNA and 1 ug Cas9 (Invitrogen) was delivered into 0.25 x 10^6^ T cells in 10 µl of buffer T, using the Neon Transfection System (Thermo Fisher Scientific). Electroporation was performed with three 1600V-10ms pulses. Gene knock-out was performed 48 hrs post T cell activation and 24 hrs prior to γ-retroviral transduction. To confirm 4-1BB KO, cells were stimulated with PMA/Ionomycin (Cell Activation Cocktail without Brefeldin A, Biolegend) for 4 hrs prior to staining for the 4-1BB protein by flow cytometry.

### Flow cytometry

For surface marker staining, cells were stained with fluorochrome-conjugated antibodies for 20 min at room temperature. For intracellular staining, cells were first stained for surface antigens, followed by intracellular staining using the BD Cytofix/Cytoperm™ Fixation/Permeabilization Kit (BD Biosciences, San Jose, CA) according to the manufacturer’s instructions. All samples were acquired on a Gallios Flow Cytometer (Beckman Coulter Life Sciences, Indianapolis, IN) or a Cytek Northern Lights (Cytek Biosciences, Inc., Fremont, CA), and data was analyzed using FlowJo 10.9.0 (BD Biosciences), Kaluza 2.4 Flow Analysis Software (Beckman Coulter Life Sciences), or Cytobank (Beckman Coulter Life Sciences) using built-in tSNE-CUDA and FlowSOM algorithms. Heatmaps were generated with hierarchical clustering using complete linkage with spearman correlation on the heatmapper software^70^. Antibodies used in this study are listed in Supplementary Table 1.

### Phospho-flow staining

Gene-modified T cells were rested overnight in 10% FBS-supplemented CTL medium without exogenous cytokines. Rested cells were then stimulated with plate-bound OKT3 in the presence of Monensin (BD GolgiStop, BD Biosciences) for 4 hrs. Cells were harvested and stained for surface antigens at room temperature for 20 min. After washing, cells were fixed with 2.5% formaldehyde solution (F1635, Sigma-Aldrich) at room temperature for 10 min, washed, permeabilized with pre-chilled 100% methanol (Fisher Scientific, Pittsburgh, PA) for 30 min on ice, and then washed three times. Cells were subsequently stained with the phospho-antibody for 45 min at room temperature. After incubation, cells were analyzed using a flow cytometer.

### Repeated antigen stimulation assay and RNA-seq sample preparation

For monitoring T cell expansion, 250,000 freshly thawed gene-modified T cells were cocultured with 125,000 irradiated K562-OKT3 cells in 1 mL 10% FBS-supplemented CTL medium in a 48-well plate. Cells were harvested every 3 days and 250,000 T cells were rechallenged with fresh 125,000 irradiated K562-OKT3 cells; cells were also collected for phenotype evaluation by flow cytometry. For RNA-seq sample preparation, unstimulated and stimulated cells at the indicated time points were sorted for CD8(+)ΔNGFR(+) T cell population on the cell sorter. Sorted cell pellets were stored at-80°C. Total RNA was extracted (RNeasy mini kit, Qiagen), quantified on a Nanodrop, and sequenced with 20 million paired-end reads/sample with 2 × 150 bp Illumina next-generation sequencing at Azenta Life Sciences.

### Bulk RNA-seq analysis

Sequence reads were trimmed using Trimmomatic v.0.36 to remove residual adaptor sequences and poor-quality reads. The trimmed reads were mapped to the reference human genome GRCh38.p14 (GCA_000001405.29) available on ENSEMBL using the STAR aligner v.2.5.2b to generate BAM files. Unique gene hit counts were calculated using feature counts from the Subread package v.1.5.2. Only unique reads that fell within exon regions were counted. Gene expression values were normalized using the variance-stabilizing transformation (vst) implemented in DESeq2 to remove dependence on experimental conditions, followed by differential gene expression (DEG) comparison between groups. Principal component analysis (PCA) was performed on the transposed vst matrix using the prcomp function in R, and the plots were generated with *ggplot2*. The proportion of variance explained by each principal component was calculated from the eigenvalues of the covariance matrix. Differential gene expression (DEG) analysis was performed by Azenta Life Sciences. Volcano plots of the DEGs were generated in GraphPad Prism 10 (GraphPad Software, La Jolla, CA, USA). Normalized gene expression values were used to generate the z-score heatmap encompassing all genes using a Python (v3.12) script. The script employed Matplotlib (v3.9.1) and Scipy (v1.16.3) libraries for hierarchical clustering across rows and columns using complete linkage with correlation-based distancing. The heatmap with the curated gene set was generated from the z-score of normalized gene expression values hierarchically clustered across columns (conditions) using complete linkage with spearman correlation on the heatmapper software. GSEA 4.4.0 with the reactome molecular signature database (C8) was used to investigate upregulation of the reactome gene sets based on DEGs between conditions.

### Cytokine quantification

The supernatant from gene-modified T cells stimulated with irradiated K562-OKT3 cells at 1:1 ratio for 24 hrs at the indicated stimulation cycle were collected and stored at-80°C. Cytokine concentrations were measured in the supernatant using the Human Luminex Discovery Assay (Merck Millipore, Billerica, MA), and samples were acquired on the Luminex 200 instrument (Thermo Fisher Scientific, Invitrogen, Grand Island, NY) according to the manufacturer’s instructions.

### Coculture experiments

For the coculture experiments, freshly thawed effector T cells were cocultured with 20,000 GFP(+) tumor cells at the indicated E:T ratio for each experiment in 200 μL of medium without exogenous cytokines in a 96-well flat-bottom plate for 9 days (4 wells/condition). One well/condition was harvested every 3 days, starting on Day 0, stained, and analyzed using a flow cytometer. To quantify cell numbers by flow cytometry, 10 μL/sample of CountBright Absolute Counting Beads (Thermo Fisher Scientific) was added, and 7-AAD (BD Biosciences) was used to exclude dead cells. Acquisition was halted at 1,000 beads and cell number was calculated.

### NFκB reporter assay

To generate the Jurkat reporter cell line, Jurkat cells were modified to express NFkB/GFP-Q8, ΔFas-ΔNGFR, and 4-1BB. The transduced population was purified using either a cell sorter or the MACS system. The Jurkat reporter cell line was then incubated with gene-modified activated T cells for 24 hrs in the presence of 500 nM Dasatinib (Selleckchem) and 100uM Z-IETD-FMK Caspase-8 Inhibitor (Selleckchem) to block T cell-mediated killing of Jurkat cells. After incubation, cells were harvested and analyzed for GFP induction within Jurkat cells on a flow cytometer.

### Microscopy sample preparation and immunofluorescence imaging

Gene-modified T cells were washed with PBS and fixed with 4% formaldehyde solution for 15 min at room temperature. After washing, cells were stained with Hoechst nucleic acid stain (Thermo Fischer Scientific) for 5 min at room temperature. Stained cells were washed, resuspended at 50,000 cells/150 uL in PBS, and deposited onto microscopic glass slides (#1.5 thickness) via spinning at 500 rpm for 5 min at room temperature in a Cytospin centrifuge (Thermo Shandon, Marshall Scientific). Samples were mounted with glycerol solution (pH: 8.5), and coverslip was placed and sealed with nail polish. Microscopic images were captured on an inverted Leica Stellaris confocal microscope (Leica Microsystems) at the Baylor College of Medicine Microscopy Core Facility using a 63X oil-emersion objective lens. Hoechst emission was captured at 461 nm wavelength, mEmerald at 509 nm and mScarlet at 594 nm. Images were analyzed using Fiji 1.0.0 software.

### Western blots

Gene-modified T cells were either left unstimulated or stimulated for 24 hrs on OKT3-coated non-tissue culture 24 well plates. After incubation, cells were harvested, and total cell lysates were prepared using RIPA lysis buffer (Sigma-Aldrich) and Halt™ Protease and Phosphatase Inhibitors (Thermo Scientific) for 30 mins on ice with intermittent vortexing. Cell lysates were passed through a fine needle to shear genomic DNA. For gel electrophoresis, lysate samples were prepared in Laemmli buffer with β-mercaptoethanol (reduced gel), incubated at 95°C for 5 min, and loaded onto 10-12% Mini-PROTEAN® TGX™ Precast Gels (Bio-Rad, Hercules, CA). Western blotting was performed by wet transfer to nitrocellulose membranes for 90 min. Membranes were blocked in 5% milk for 1hr at room temperature, followed by incubation with primary antibodies diluted in 5% milk overnight at 4°C with shaking. Next day, membranes were washed three times and incubated with the corresponding luminescent secondary antibodies for 1 hr at room temperature with shaking before visualization on the LI-COR Odyssey imaging system (LI-COR Biotechnology, Lincoln, NE). Antibodies used for western blot are listed in Supplementary Table 1.

### Quantitative RT-PCR

Total RNA was extracted (RNeasy mini kit, Qiagen) from gene-modified T cells that were either unstimulated or stimulated for 24 hrs on OKT-3-coated plates. RNA was reverse transcribed into cDNA using the Superscript IV First-Strand Synthesis System kit (Thermo Fisher Scientific) according to the manufacturer’s instructions. qPCR was performed in triplicate on the cDNA template using the 4-1BB-specific forward primer (ACTGCCCAGCTGGTACATTC) and reverse primer (TTGCTGGTGGAGGAACACTC), generating a 160-bp amplicon. β-Actin was used as an internal control with the forward primer (AGAGCTACGAGCTGCCTGAC) and reverse primer (GGATGCCACAGGACTCCA), generating a 111-bp amplicon. Reactions were performed using iTag Universal SYBR Green Supermix (Bio-Rad) on the Bio-Rad CFX96 System.

### In vivo studies

Breeder pairs of NOD.Cg-*Prkdc^scid^* H2-K1*^b-tm1Bpe^* H2-Ab1*^g^*^7^*^-em1Mvw^* H2-D1*^b-tm1Bpe^* Il2rg^tm1Wjl^/SzJ mice (NSG-MHC I/II DKO mice; Strain no. 025216) were purchased from the Jackson Laboratory and bred in the Baylor College of Medicine animal facility. Both female and male littermates (8-12 weeks) were used for experiments. All animal experiments were conducted in compliance with the Baylor College of Medicine Institutional Animal Care and Use Committee (IACUC; protocol no. AN-4758). To evaluate the effect of DSR on the *in vivo* anti-tumor efficacy of Survivin TCR-modified T cells, 3 x 10^6^ BV173-CBG/GFP cells were first injected intravenously. Seven days later, freshly thawed 5 x 10^6^ AkaLuc(+) Survivin TCR(+) CD8(+) T cells were injected intravenously. For the T-cell engager model, mice were injected intravenously with 0.5 x 10^6^ MOLM-13-CBG/GFP cells followed 4 days later by the intravenous injection of freshly thawed 5 x 10^6^ AkaLuc(+) T cells (comprising 60-70% T-cell engager(+) T cells). For the cTCR model, 0.5 x 10^6^ NALM6-CBG/GFP cells were injected intravenously, and 7 days later, freshly thawed 0.5 x 10^6^ AkaLuc(+) cTCR(+) T cells were injected intravenously. In all animal experiments, both tumor cell growth and T cell expansion/persistence were tracked using a dual-luciferase bioluminescence imaging system optimized in our laboratory. Briefly, mice were injected intraperitoneally with 100 μL TokeOni (1 mM; Sigma-Aldrich) and imaged for AkaLuc bioluminescence using an IVIS Lumina III imaging system (Caliper Life Sciences, Hopkinton, MA, USA). Subsequently, mice were injected with 100 μL D-luciferin (30 mg/mL; PerkinElmer, Waltham, MA, USA) and imaged with a 520-nm filter for CBG bioluminescence. Data was analyzed using the Living Image software (Caliper Life Sciences). To evaluate the surface phenotype of T cells, a total of 100 ul peripheral blood was collected once in two weeks from the tail vein for flow cytometry analysis.

## Statistical analysis

Statistical analysis was performed using GraphPad Prism 10 and 11. The statistical tests applied in each experiment are described in the corresponding figure legends.

## Acknowledgements

We thank Dr. Bilal Omer at the Baylor College of Medicine for kindly gifting the γ-retroviral vector for Survivin TCR. Bulk RNA sequencing and data pre-processing was performed at Azenta Life Sciences. We thank Roopal Garg at Google LLC for contributing to the generation of the heatmap for complete gene expression dataset. We also thank Dr. Chiou-Tsun Tsai for assistance in western blot experiments. We would like to acknowledge the Baylor College of Medicine small animal core and imaging facility for maintaining the animal breeding colonies and for giving access to the IVIS equipment. Thanks to the Baylor College of Medicine microscopy core for training and access to the Leica Stellaris confocal imaging microscope. Images were created in BioRender. Narula, M. (2026) https://BioRender.com/w6vf8j7.

## Author contributions

NW conceptualized and designed the study. MN, JE and NW developed the methodology. MN, JE, CO, TH, AA, FM and NW performed the experiments. MN and NW analyzed the data and interpreted the results. ISS performed the computational analysis. MN and NW wrote the original draft of the manuscript. MM secured the funding. NW supervised and managed project administration. MN, NW and MM reviewed and edited the manuscript.

## Data Availability

All data and materials used to complete and draw conclusions in this study are presented in the paper, including supplementary information.

