## Supplementary Figures for "Integrating activation-induced costimulation and cytokine signals enhance TCR-based cell therapies"

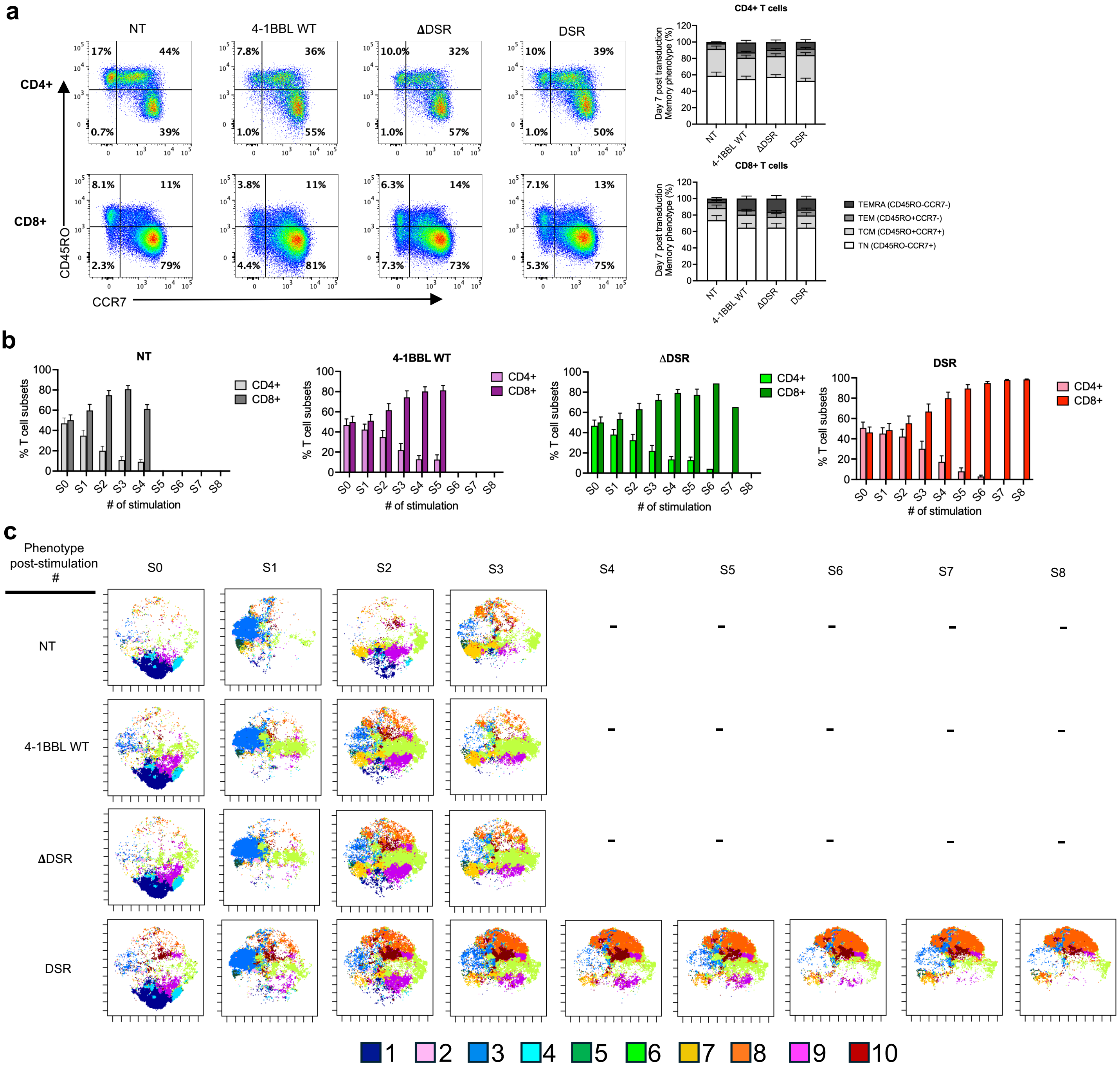


**Extended Figure 1: Characterization of gene-modified T cells before and after repeated antigen stimulation. a,** Representative flow plots and summary stacked bar graphs of the memory phenotype (CD45RO/CCR7) in CD4(+) (top) and CD8(+) (bottom) T cell subsets on day 7 post-transduction (n=12). **b,** Distribution of CD4(+)/CD8(+) T cell subsets within each experimental group at the end of each stimulation cycle of the repeated antigen stimulation (n=9). **c,** tSNE-CUDA plots of the cell phenotype showing the distribution of 10 metaclusters within each experimental group and each stimulation cycle of the repeated antigen stimulation (n=5). All data with multiple donors shown as Mean ± S.E.

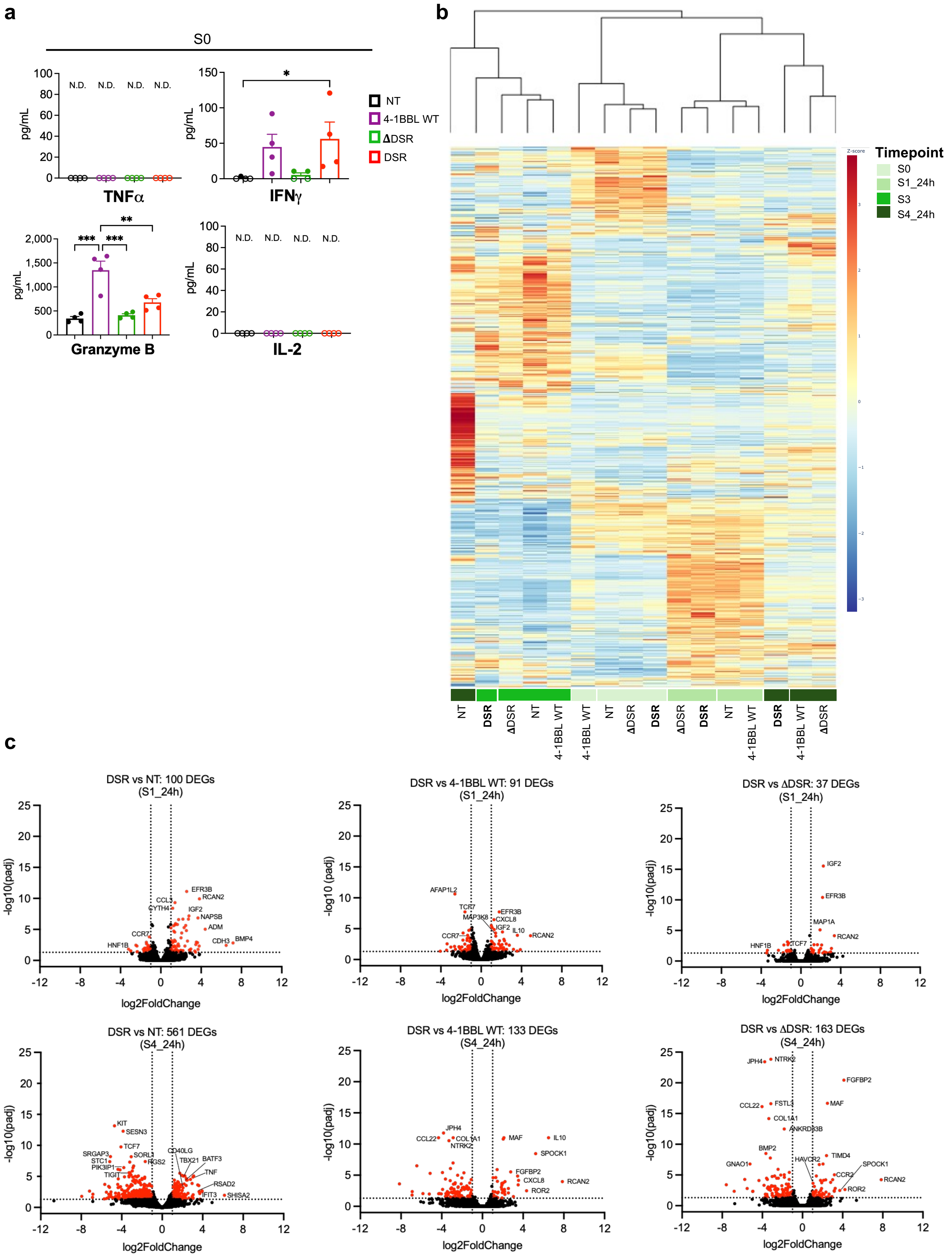


**Extended Figure 2**

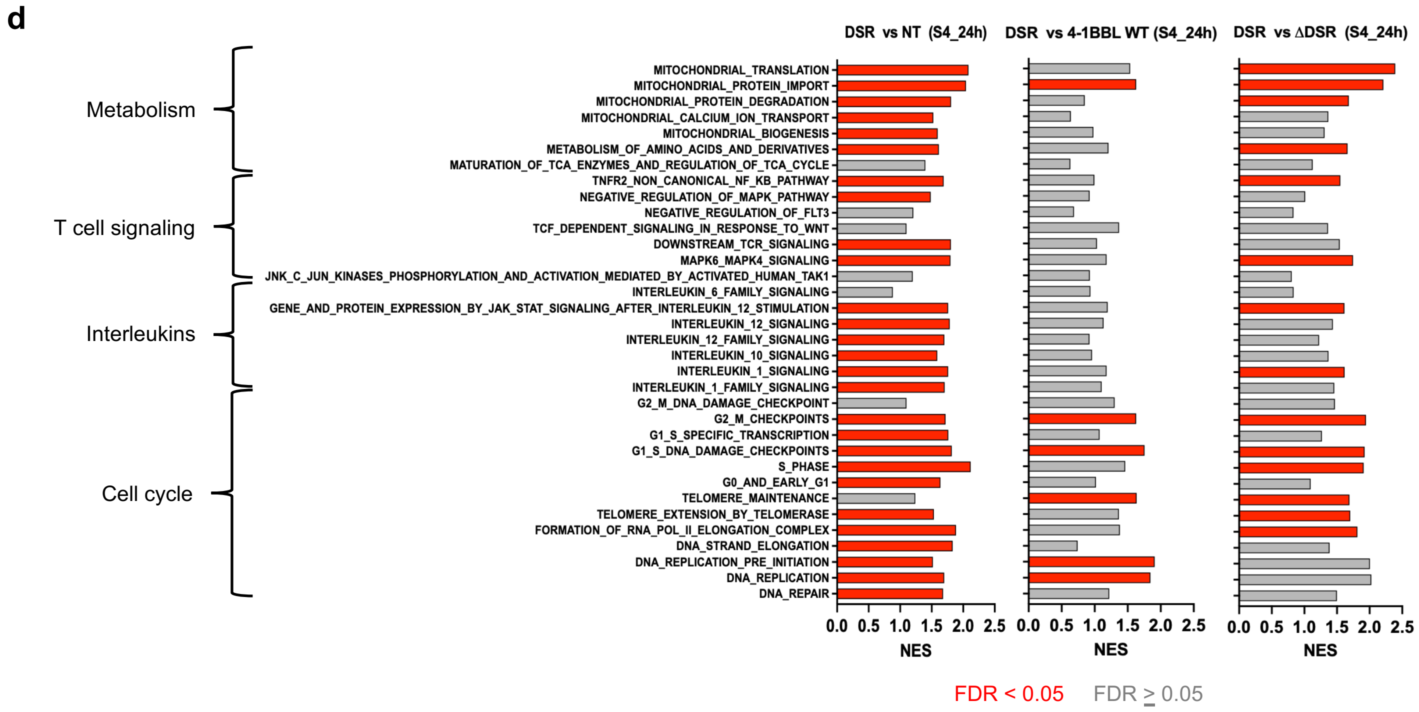


**Extended Figure 2: Transcriptome comparison of DSR-modified T cells versus control groups upon repeated antigen stimulation. a,** Summary plots of effector cytokine secretion in unstimulated (S0) T cells (Mean ± S.E, n=4). **b,** z-score heatmap of the normalized gene expression (mean of 3 donors) of the entire gene set (~17,000 genes) across groups and time points specified in Figure 2a of the repeated stimulation assay with hierarchical clustering across rows. **c,** Volcano plots showing differentially expressed genes (DEGs in red) in DSR-modified T cells compared to each control group (NT, 4-1BBL WT, ΔDSR) at S1_24h (top) and S4_24h (bottom). Dotted lines show the DEG cut-off at -log10(padj) value of ≥1.3 and log2 Fold Change ≥ |1|. **d,** GSEA Reactome pathways common to and upregulated in DSR-modified T cells compared to each control group (NT, 4-1BBL WT, ΔDSR); significantly upregulated pathways indicated in red. Statistical analysis was performed by RM one-way ANOVA with Tukey's multiple comparison in panel a. *p ≤ 0.05, **p ≤ 0.01, ***p ≤ 0.001, ****p ≤ 0.0001. N.D. Not detectable, GSEA: Gene-set enrichment analysis.

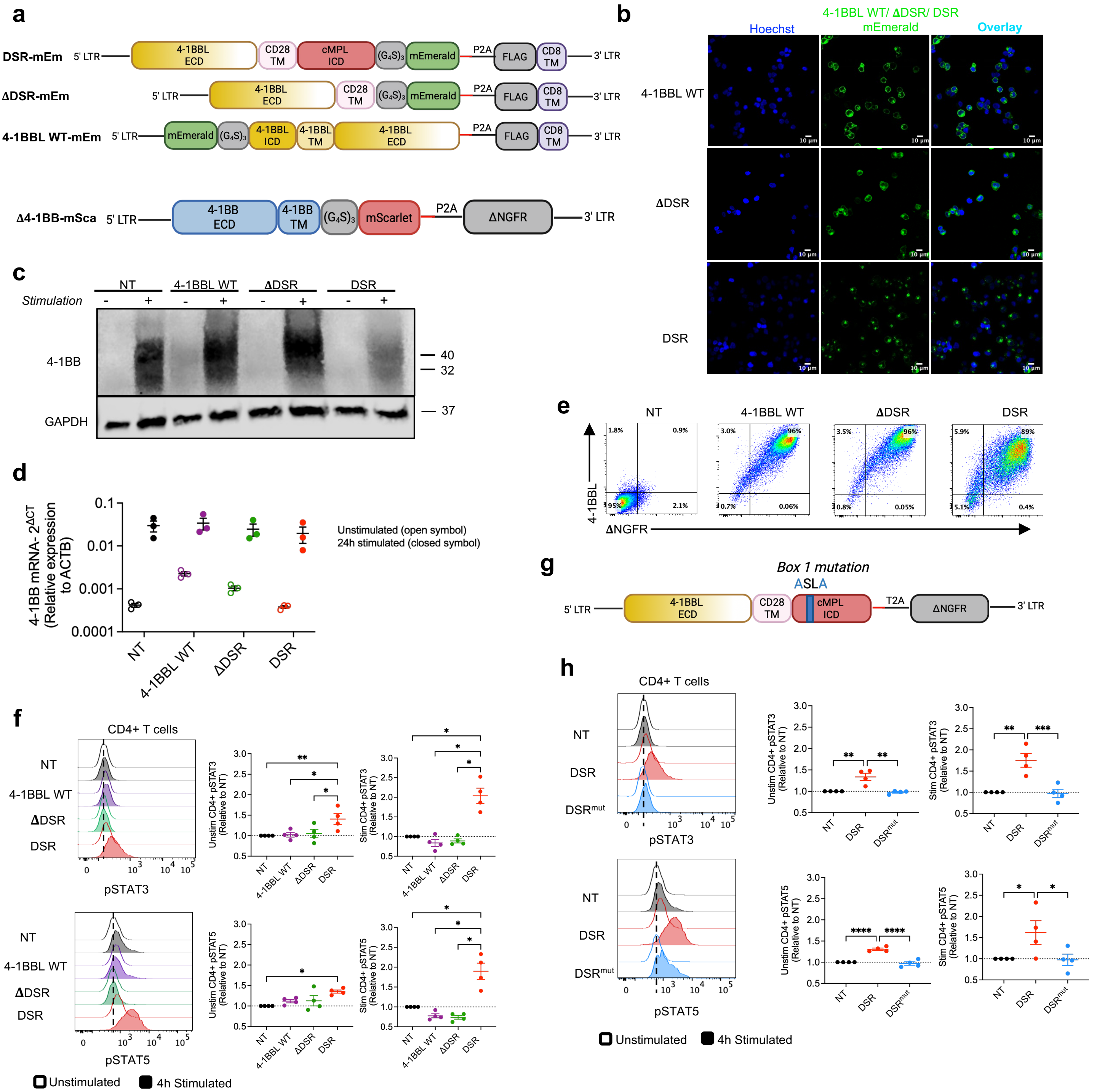

**Extended Figure 3: DSR interaction with 4-1BB. a,** Bicistronic γ-retroviral construct maps of mEmerald-tagged DSR, ΔDSR and 4-1BBL WT with FLAG surrogate marker (top) and mScarlet-tagged Δ4-1BB with ΔNGFR surrogate marker (bottom). **b,** Fluorescence microscopy images of T cells gene-modified with mEmerald (green)-tagged 4-1BBL WT, ΔDSR or DSR. Hoechst (blue) indicates nuclear stain. Images were acquired with a 63x objective lens. **c,** Western blot image showing total 4-1BB protein (32 and 40 kDa bands) in T cells with (+) or without (-) 24 hr OKT3-stimulation. GAPDH (37kDa) was used as an internal control. **d,** 4-1BB mRNA expression relative to β-actin housekeeping gene by quantitative PCR in unstimulated (open circle) and 24 hr OKT3-stimulated (closed circle) T cells (n=3). **e,** Flow plots of 4-1BBL surface expression in gene-modified T cells cocultured with Jurkat cells (48 hr post stimulation). **f,** Representative histograms of phospho-STAT3 and phospho-STAT5 staining in unstimulated (open) or 4 hr OKT3-stimulated (filled) CD4(+) T cells (left). Summary plots showing gMFI relative to NT cells (n=4, right). **g,** γ-retroviral construct map of DSR mutant with mutation in the Box1-JAK2 binding site within the intracellular (ICD) domain. **h,** Representative histograms of phospho-STAT3 and phospho-STAT5 staining in unstimulated (open) or 4 hr OKT3-stimulated (filled) CD4(+) T cells modified with DSR mutant (DSR$^{mut}$). Summary plots showing gMFI relative to NT cells (n=4, right).

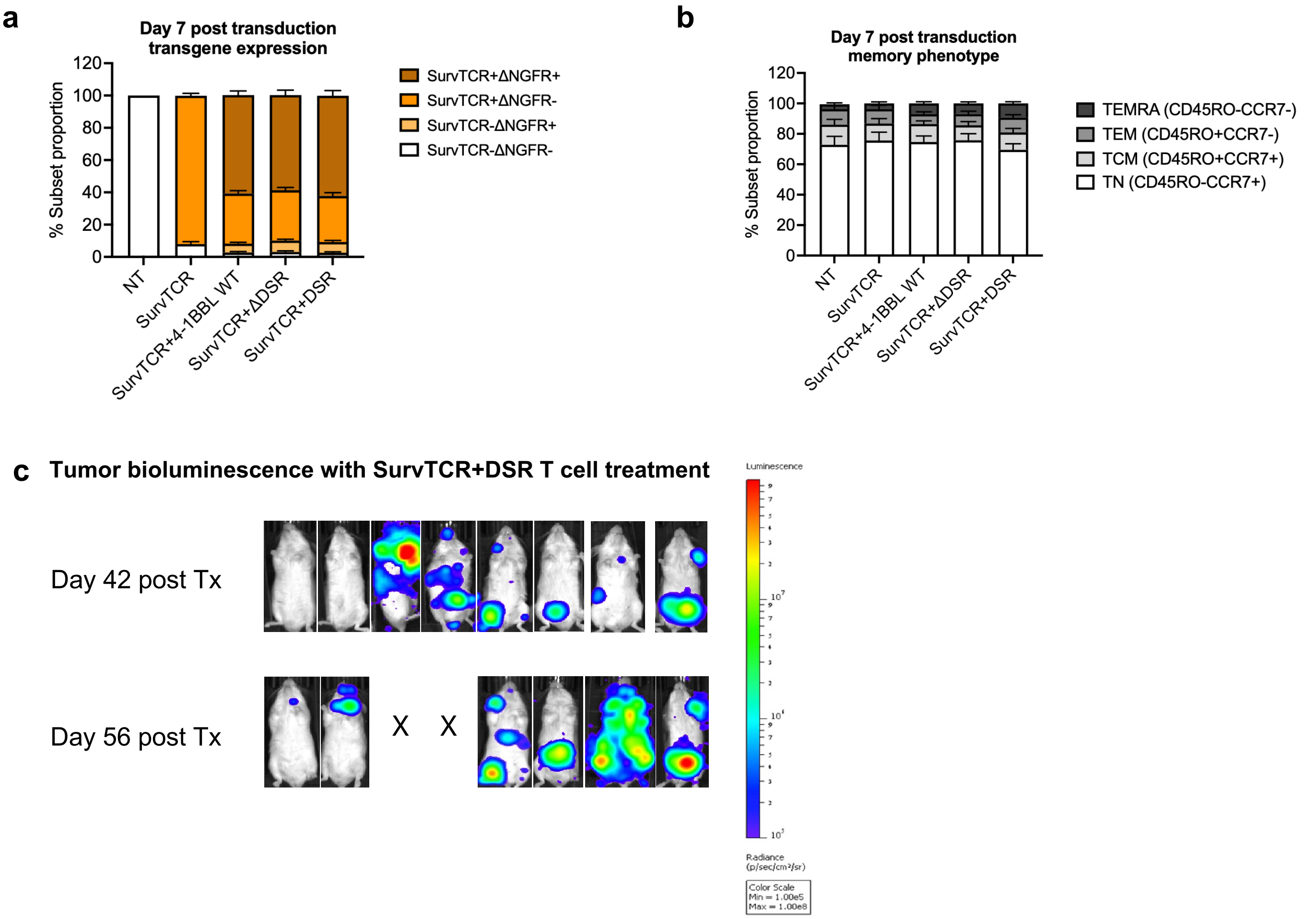


**Extended Figure 4: Characterization of DSR-engineered Survivin TCR-T cells.** Summary stacked bar graphs showing the transgene expression (Survivin TCR and ΔNGFR - as a surrogate marker for 4-1BBL WT/ΔDSR/DSR) (**a**), and memory phenotype (CD45RO/CCR7) (**b**) of the gene-modified CD8(+) T cell products (day 7 post transduction, Mean ± S.E., n=6). **c,** Tumor bioluminescence on day 42 (top) and day 56 (bottom) post treatment (Tx) with DSR-modified Survivin TCR-T cells (n=8).

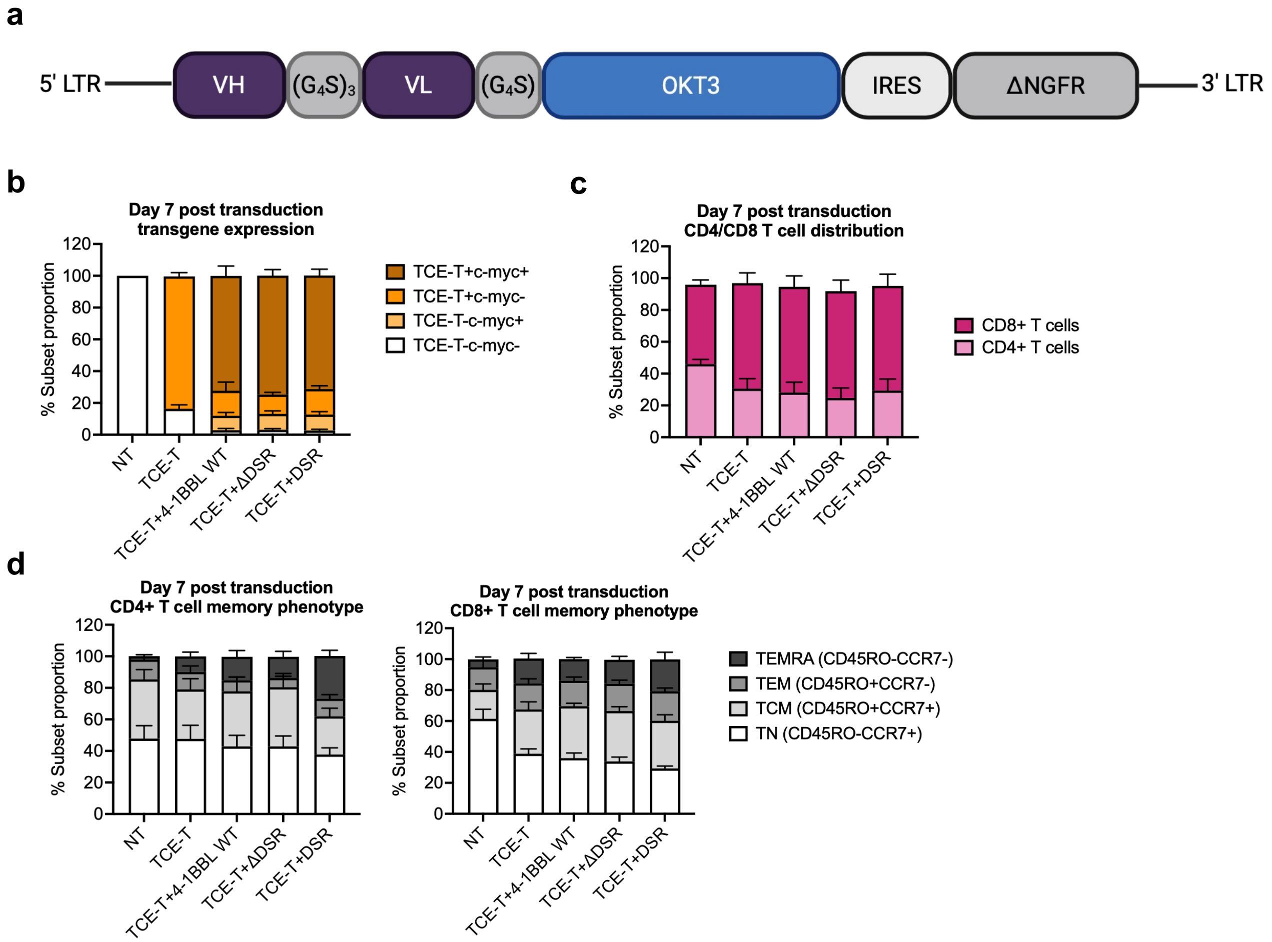


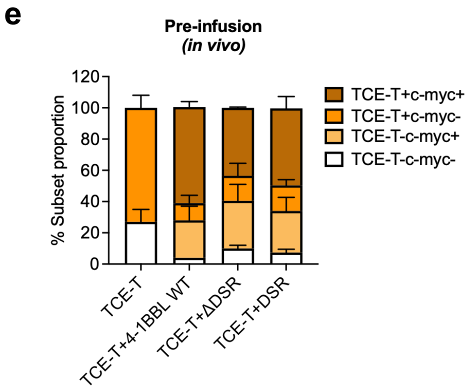


**Extended Figure 5: Characterization of T cell engager secreting-T cells (TCE-T). a,** γ-retroviral construct map of the bicistronic CD123-targeting TCE. **b-d,** Summary stacked bar graphs showing the phenotype of gene-modified T cells (day 7 post transduction, n=7); transgene expression (TCE-T - ΔNGFR, 4-1BBL WT/ΔDSR/DSR - c-myc as the surrogate marker) (**b**), CD4(+)/CD8(+) T cell distribution (**c**), memory phenotype (CD45RO/CCR7) of CD4(+) T cells (left) and CD8(+) T cells (right) (**d**). **e,** Summary stacked bar graphs showing the transgene expression of infused T cell products (TCE-T - ΔNGFR, 4-1BBL WT/ΔDSR/DSR - c-myc as the surrogate marker) (day 7 post transduction, n=2). Data with multiple donors shown as Mean ± S.E for b,c,d,e.

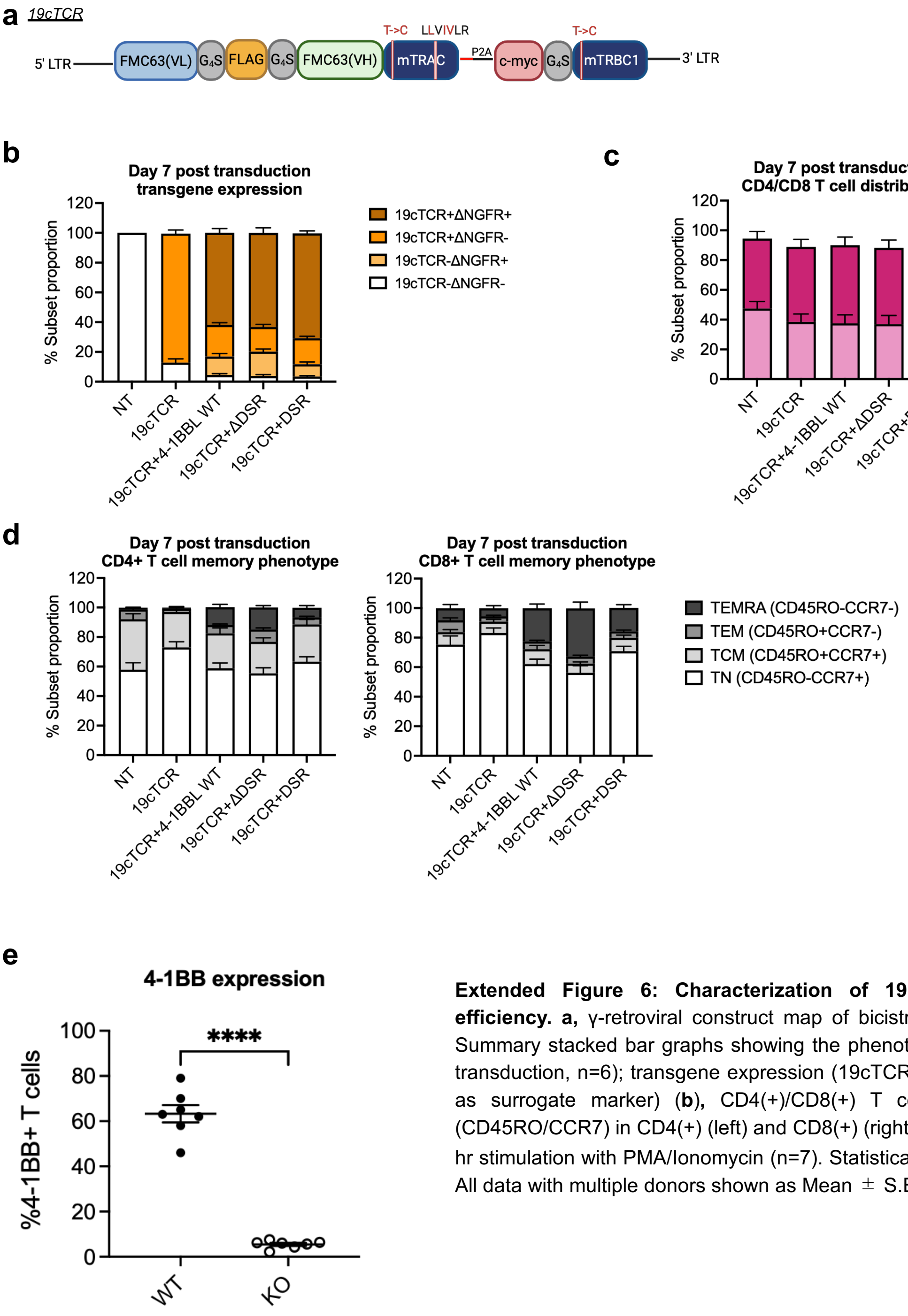


**Extended Figure 6: Characterization of 19cTCR-T cells and 4-1BB knock-out efficiency. a,** γ-retroviral construct map of bicistronic CD19-targeting chimeric TCR. **b-d,** Summary stacked bar graphs showing the phenotype of gene-modified T cells (day 7 post transduction, n=6); transgene expression (19cTCR-FLAG, 4-1BBL WT/ΔDSR/DSR- ΔNGFR as surrogate marker) (**b**), CD4(+)/CD8(+) T cell distribution (**c**), memory phenotype (CD45RO/CCR7) in CD4(+) (left) and CD8(+) (right) (**d**). **e,** 4-1BB surface expression upon 4 hr stimulation with PMA/Ionomycin (n=7). Statistical analysis was performed by paired t-test. All data with multiple donors shown as Mean ± S.E.. WT: wild-type, KO: knock-out
